# Protein polyglutamylation catalyzed by the bacterial Calmodulin-dependent pseudokinase SidJ

**DOI:** 10.1101/738567

**Authors:** Alan Sulpizio, Marena E. Minelli, Min Wan, Paul D. Burrowes, Xiaochun Wu, Ethan Sanford, Jung-Ho Shin, Byron Williams, Michael Goldberg, Marcus B. Smolka, Yuxin Mao

## Abstract

Pseudokinases are considered to be the inactive counterparts of conventional protein kinases and comprise approximately 10% of the human and mouse kinomes. Here we report the crystal structure of the *Legionella pneumophila* effector protein, SidJ, in complex with the eukaryotic Ca^2+^-binding regulator, Calmodulin (CaM). The structure reveals that SidJ contains a protein kinase-like fold domain, which retains a majority of the characteristic kinase catalytic motifs. However, SidJ fails to demonstrate kinase activity. Instead, mass spectrometry and in vitro biochemical analysis demonstrate that SidJ modifies another *Legionella* effector SdeA, an unconventional phosphoribosyl ubiquitin ligase, by adding glutamate molecules to a specific residue of SdeA in a CaM-dependent manner. Furthermore, we show that SidJ-mediated polyglutamylation suppresses the ADP-ribosylation activity. Our work further implies that some pseudokinases may possess ATP-dependent activities other than conventional phosphorylation.

## Introduction

Phosphorylation mediated by protein kinases is a pivotal posttranslational modification (PTM) strategy affecting essentially every biological processes in eukaryotic cells (Brognard and Hunter, 2011; Cohen, 2002). The importance of protein phosphorylation is further endorsed by the fact that the mammalian genome contains more than 500 protein kinases, corresponding to ∼2% of the total proteins encoded in the genome (Manning et al., 2002; Rubin et al., 2000). Despite the importance of phosphorylation, about 10% of kinases of the mammalian kinome lack key catalytic residues and are considered pseudokinases (Jacobsen and Murphy, 2017; Shaw et al., 2014). Accumulated evidence demonstrated that catalytically inactive pseudokinases have important noncatalytic functions, such as allosteric regulators (Scheeff et al., 2009; Zeqiraj et al., 2009) or nucleation hubs for signaling complexes (Brennan et al., 2011; Jagemann et al., 2008). Interestingly, a recent study uncovered AMPylation activity catalyzed by an evolutionary conserved pseudokinase selenoprotein (SelO) (Sreelatha et al., 2018). The SelO pseudokinases bind ATP with a flipped orientation relative to the ATP bound in the active site of canonical kinases and transfer the AMP moiety, instead of the γ-phosphate, from ATP to Ser, Thr, or Tyr residues on protein substrates. This finding suggests that pseudokinases should be reconsidered for alternative ATP-dependent PTM activities.

Protein glutamylation is another type of ATP-dependent PTM, in which the γ-carboxyl group of a glutamate residue in a targeted protein is activated by ATP and then forms a isopeptide bond with the amino group of a free glutamate. Alternatively, multiple glutamates can be sequentially added to the first to generate a polyglutamate chain (Janke et al., 2008). Protein glutamylation was first discovered on the proteins that build microtubules, the α-tubulins and β-tubulins (Alexander et al., 1991; Edde et al., 1990; Redeker et al., 1992; Rudiger et al., 1992). Further studies revealed that tubulin polyglutamylation is mediated by a group of tubulin tyrosine ligase-like (TTLL) family glutamylases (van Dijk et al., 2007). These glutamylases belong to ATP-grasp superfamily and have a characteristic fold of two α/β domains with the ATP-binding active site situated between them (Garnham et al., 2015; Szyk et al., 2011). So far, the TTLL polyglutamylases are the only family of enzymes catalyzing protein glutamylation although new polyglutamylated substrates have been identified besides tubulins (van Dijk et al., 2008).

The facultative intracellular pathogen *Legionella pneumophila* is the causative agent of Legionnaires’ disease, a potentially fatal pneumonia (McDade et al., 1977; McKinney et al., 1981). *L. pneumophila* delivers a large number (>300) of effector proteins into the host cytoplasm through its Dot/Icm type IV secretion system (Segal et al., 1998; Vogel et al., 1998), leading to the creation of a specialized membrane-bound organelle, the *Legionella*-containing vacuole (LCV) (Hubber and Roy, 2010; Isberg et al., 2009; Lifshitz et al., 2013; Zhu et al., 2011). Among the large cohort of *Legionella* effectors, the SidE family of effectors have recently been identified as a group of novel Ub ligases that act independently of ATP, Mg^2+^ or E1 and E2 enzymes (Bhogaraju et al., 2016; Kotewicz et al., 2017; Qiu et al., 2016). This unusual SidE family ubiquitin ligases contain multiple domains including a mono-ADP-ribosyl transferase (mART) domain, which catalyzes ubiquitin ADP-ribosylation to generate mono-ADP-ribosyl ubiquitin (ADPR-Ub), and a phosphodiesterase (PDE) domain, which conjugates ADPR-Ub to serine residues on substrate proteins (phosphoribosyl-ubiquitination) (Akturk et al., 2018; Dong et al., 2018; Kalayil et al., 2018; Kim et al., 2018; Wang et al., 2018). Interestingly, the function of SidEs appears to be antagonized by SidJ (Lpg2155), an effector encoded by a gene resides at the same locus with genes encoding three members of the SidE family (Lpg2153, Lpg2156, and Lpg2157) (Liu and Luo, 2007). It has been shown that SidJ suppresses the yeast toxicity conferred by the SidE family effectors (Havey and Roy, 2015; Jeong et al., 2015; Urbanus et al., 2016). Furthermore, SidJ has been shown to act on SidE proteins and releases these effectors from the LCV (Jeong et al., 2015). A recent study reported that SidJ functions as a unique deubiquitinase that counteracts the SidE-mediated phosphoribosyl-ubiquitination by deconjugating phosphoribosyl-ubiquitin from modified proteins (Qiu et al., 2017). However, our recent results do not support this SidJ-mediated deubiquitinase activity (Wan et al., 2019) and the exact function of SidJ remains elusive.

The goal of the present study is to elucidate the molecular function of SidJ and to investigate the mechanism that underlies how SidJ antagonizes the PR-ubiquitination activity of SidEs. Here we report the crystal structure of SidJ in complex with human Calmodulin 2 (CaM) and reveal that SidJ adopts a protein kinase-like fold. A structural comparison allowed us to identify all the catalytic motifs conserved in protein kinases. However, SidJ failed to demonstrate protein kinase activity. Using SILAC (Stable Isotope Labeling by Amino acids in Cell culture) based mass spectrometry approach, we discovered that SidJ modifies SdeA by attaching the amino acid glutamate to a key catalytic residue on SdeA. Moreover, we found that this glutamylation activity by SidJ is CaM dependent and the glutamylation of SdeA suppresses its PR-ubiquitination activity. Thus our work provides molecular insights of a key PR-ubiquitination regulator in *Legionella* infection. We anticipate that our work will also have impact on the studies of pseudokinases and CaM-regulated cellular processes.

## Results

### SidJ Binds CaM through its C-terminal IQ Motif

To elucidate the biological function of SidJ, we performed sequence analyses and found that the C-terminus of SidJ contains the sequence “IQxxxRxxRK”, which resembles the IQ motif found in a number proteins, mediating the binding with Calmodulin (CaM) in the absence of Ca^2+^ (Figure 1A) (Rhoads and Friedberg, 1997). To test whether this predicted IQ motif in SidJ can mediate an interaction with CaM, we prepared recombinant proteins of SidJ and CaM and incubated these proteins in the presence or absence of Ca^2+^. We then analyzed the samples with native PAGE and observed that a new band corresponding to the SidJ-CaM complex appeared in a Ca^2+^ independent manner (Figure 1B). The formation of the complex was dependent on the intact IQ motif as the SidJ IQ mutant (I841D/Q842A) did not form a stable complex with CaM. The interaction between SidJ and CaM was further quantified by isothermal calorimetry (ITC) analysis, which showed a dissociation constant (Kd) of about 89.6 nM between CaM and wild type SidJ with a 1:1 stoichiometry, while no binding was detected between CaM and SidJ IQ mutant (Figure 1C). The association between SidJ and CaM was also demonstrated by size exclusion chromatography as the wild type SidJ and CaM co-fractionated while the SidJ IQ mutant migrated separately from CaM (Figure 1—figure supplement 1). Collectively, SidJ interacts with CaM through its C-terminal IQ motif in a Ca^2+^ independent manner.

**Figure 1.**
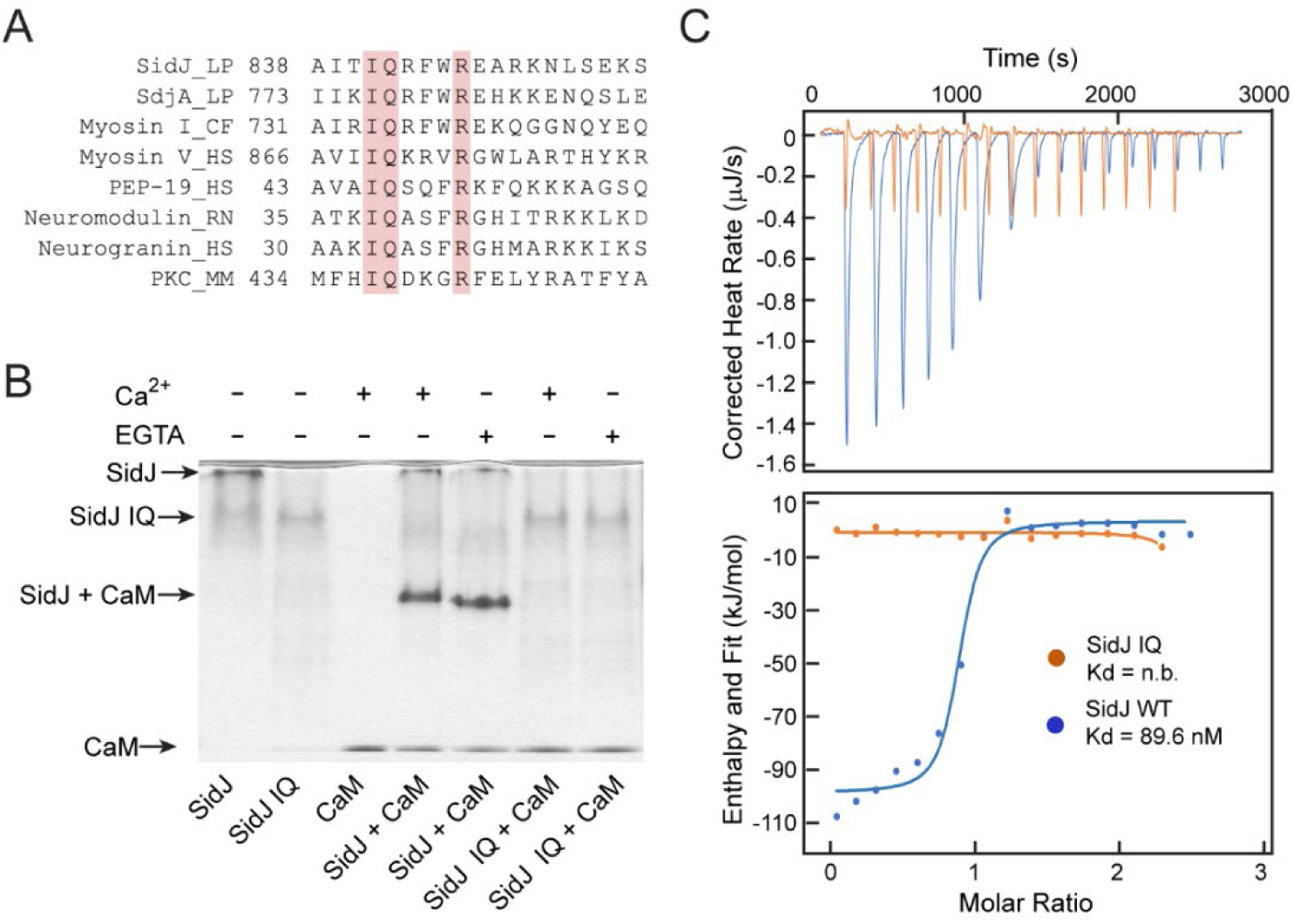
SidJ Binds CaM through its C-terminal IQ Motif. (A) Multiple sequence alignment of IQ motifs (“IQXXXR”) mediating the binding of CaM from the indicated proteins. Protein names are listed followed by a two-letter representation of the species and the residue numbers of the first amino acid in the aligned sequences. Identical residues of the motif are highlighted in salmon. Entrez database accession numbers are as follows: SidJ, YP_096168.1; SdjA, YP_096515.1; Myosin-1, ONH68659.1; Myosin V, NP_000250.3; PEP-19, CAA63724.1; Neuromodulin, NP_058891.1; Neurogranin, NP_006167.1; Protein kinase C delta isoform, NP_001297611.1. LP: *Legionella pneumophila*; CF: *Cyberlindnera fabianii*; HS: *Homo sapiens*; RN: *Rattus norvegicus*; MM: *Mus musculus*. (B) Native PAGE analysis of the SidJ and CaM complex. Wild type SidJ and CaM form a complex independent of Ca^2+^ and the complex migrates at a different position from each individual protein. (C) Isothermal titration calorimetry measurement of the affinity between CaM and SidJ WT (blue) or SidJ IQ mutant (orange). The top panel shows the reconstructed thermogram, and the bottom panel the isotherms. SidJ WT binding to CaM has a dissociation constant of approximately 89.6 nM in a 1:1 stoichiometry.

### Overall Structure of the SidJ and CaM Complex

Despite extensive trials, we were unable to obtain protein crystals for SidJ alone. However, the stable interaction between SidJ and CaM allowed us to crystallize SidJ in complex with CaM. The structure was determined by selenomethionine single wavelength anomalous dispersion (SAD) method and was refined to a resolution of 2.6 Å with good crystallographic R-factors and stereochemistry (Table 1). Based on the SidJ-CaM structure, the SidJ protein is comprised of four functional units: a N-terminal regulatory domain (NRD), a base domain (BD), a kinase-like catalytic domain, and a C-terminal domain (CTD) containing the CaM-binding IQ motif (Figure 2). The N-terminal portion of the NRD (residues 1-88) is predicted to be intrinsically disordered and thus was not included in the SidJ construct for crystallization trials. The rest of the NRD (residues 89-133) adopts an extended structure with three β-strands and flexible connecting loops and meanders on the surface across the entire length of the kinase-like domain (Figure 2B and 2C). The BD is mainly comprised of α-helices. It interacts with both the kinase-like domain and CTD and provides a support for these two domains to maintain their relative orientation. The CTD contains four α-helices with the first three α-helices forming a tri-helix bundle and the fourth IQ motif-containing α-helix (IQ-helix) extending away from the bundle to engage in interactions with CaM. In the SidJ-CaM complex, CaM “grips” the IQ-helix with its C-lobe (Figure 2B and 2C, left panels) while its N-lobe interacts with the NRD, CTD, and the kinase-like domains. In agreement with our biochemical results that CaM binds SidJ in a Ca^2+^ independent manner (Figure 1). Only the first EF-hand of CaM is observed to coordinate with a Ca^2+^ ion based on the difference Fourier electron density map even though the crystal is formed in a crystallization buffer containing 1 mM CaCl_2_ (Figure 2—figure supplement 1).

**Figure 2.**
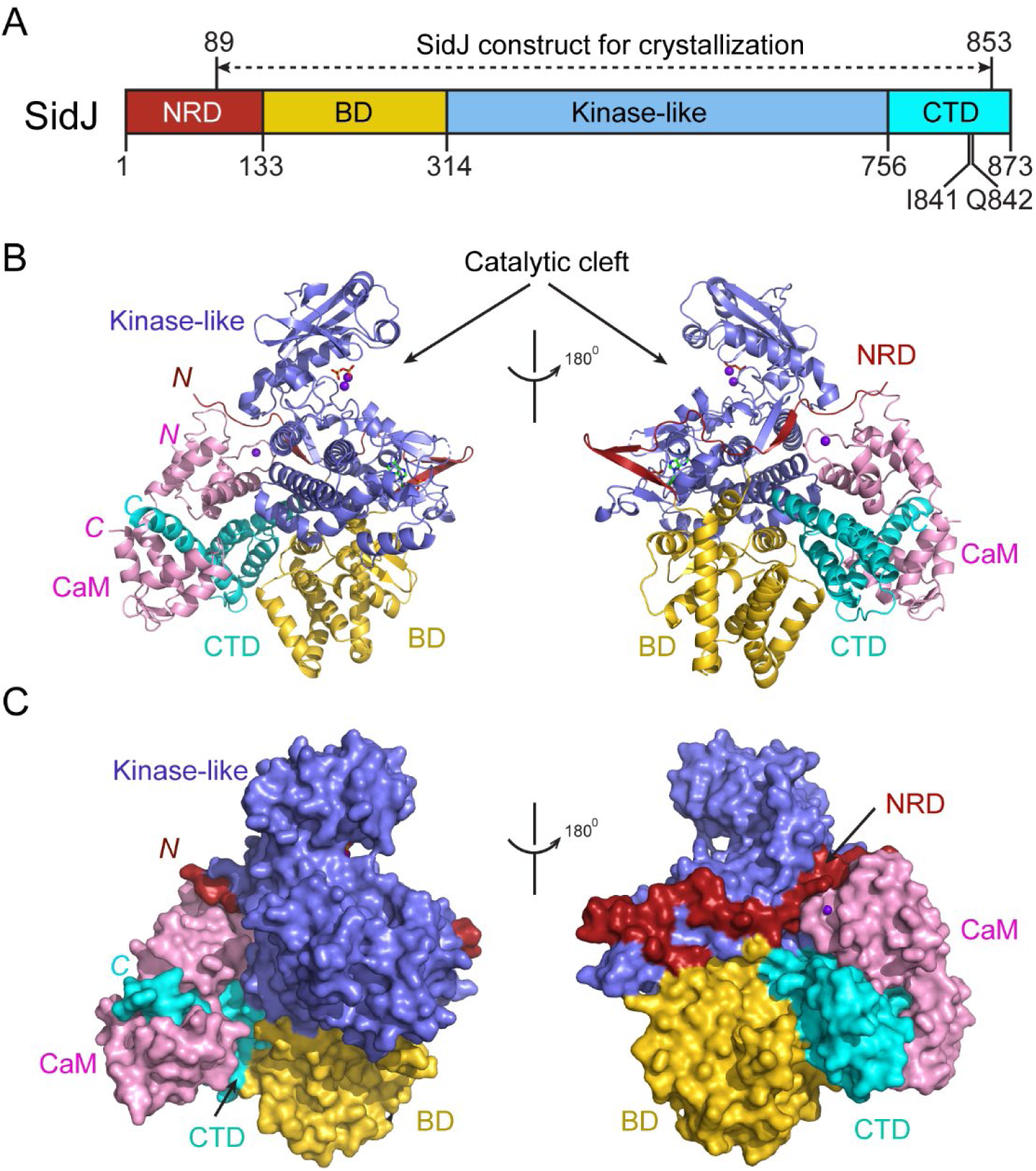
Overall structure of the SidJ and CaM complex. (A) Schematic diagram of SidJ domain architecture. SidJ is comprised of a N-terminal regulatory domain (NRD) in maroon, a Base domain (BD) in yellow, kinase-like catalytic domain in blue, and a C-terminal domain (CTD) in cyan. The construct used for crystallography (89-853) is depicted above the schematic. (B) Overall structure of SidJ bound to CaM in a cartoon representation. SidJ structure is colored with the same scheme as in (A) and CaM is colored in pink. Ca^2+^ ions are depicted as purple spheres. The kinase-like domain of SidJ has a bilobed structure with two Ca^2+^ ions and a pyrophosphate molecule bound at the catalytic cleft between the two lobes. Right panel: 180° rotation of left panel and depicts the NRD domain contacts with CaM. (C) Molecular surface representation of SidJ bound to CaM in the same orientation and coloring as in (B). Right panel: 180° rotation of the left panel. Note that the NRD meanders on the surface of the kinase-like domain and mediates the contact between the kinase-like domain and CaM.

**Table 1.** Data Collection, Phasing, and Structural Refinement Statistics.

|  | SeMet SidJ-CaM | Native SidJ-CaM<br>(PDB ID: 6PLM ) |
| --- | --- | --- |
| Synchrotron beam lines | NSLS II 17-ID-1 (AMX) | NSLS II 17-ID-1 (AMX) |
| Wavelength (Å) | 0.97949 | 0.97949 |
| Space group | P2<sub>1</sub> | P2<sub>1</sub> |
| Cell dimensions |  |  |
| *a*, *b*, *c* (Å) | 105.08, 104.08, 109.65 | 105.35, 103.79, 110.19 |
| *α*, *β*, *γ* (°) | 90, 104.49, 90 | 90, 104.69, 90 |
| Maximum resolution (Å) | 2.85 | 2.59 |
| Observed reflections | 371,678 | 482,266 |
| Unique reflections | 69,809 | 69,809 |
| Completeness (%)<sup>a</sup> | 99.5 | 97.7 |
| *<I>/<σ>*<sup>a</sup> | 43.20 (15.30) | 38.20 (13.20) |
| R<sub>sym</sub><sup>a,b</sup> (%) | 0.024 (0.068) | 0.043(0.091) |
| Phasing methods | SAD | Native |
| Heavy atom type | Se | - |
| Number of heavy atoms/ASU | 12 | - |
| Resolution (Å)<sup>a</sup> | - | 29.32(2.59) |
| R<sub>crys</sub> / R<sub>free</sub> (%)<sup>a,c</sup> | - | 17.6/24.1 |
| Rms bond length (Å) | - | 0.0142 |
| Rms bond angles (°) | - | 1.8174 |
| Most favored/allowed (%) | - | 96.65/3.35 |
| Generous/Disallowed (%) | - | 0 |

<sup>a</sup> Values in parentheses are for highest-resolution shell.
<sup>b</sup>R<sub>sym</sub> = $\sum_h \sum_i |I_i(h) - \langle I(h) \rangle| / \sum_h \sum_i I_i(h)$ .
<sup>c</sup>R<sub>crys</sub> = $\Sigma(|F_{obs}| - k|F_{cal}|) / \Sigma|F_{obs}|$ . R<sub>free</sub> was calculated for 5% of reflections randomly excluded from the refinement.

### The Core of SidJ Adopts a Protein Kinase Fold

Although there is no detectable primary sequence homology to any known protein kinase, a structure homology search with the Dali server (Holm and Laakso, 2016) showed that the core of SidJ most closely resembles the Haspin kinase (Villa et al., 2009) with a Z-score of 10.1. The SidJ core, thus named kinase-like domain, assumes a classical bilobed protein kinase fold (Figure 3A-B). A detailed structural analysis revealed that the N-lobe of the SidJ kinase-like domain contains all the structural scaffolding elements conserved in protein kinases, including a 5-stranded antiparallel β-sheet and the αC helix (the secondary structural elements are named according to PKA nomenclature) (Figure 3C). Furthermore, one of the key catalytic residues, K367 in the β3 strand, is conserved among all SidJ homologs (Figure 3—figure supplement 1 and 2). This residue is positioned towards the catalytic cleft to interact with the phosphate groups of ATP for catalysis. Similar to protein kinases, this invariable Lys is coupled by a conserved Glu (E381) in the αC helix (Figure 3C). However, the “glycine-rich loop” connecting the β1and the β2 strands forms a type I β-turn structure whereas in canonical protein kinases, the corresponding loop is much longer and packs on top of the ATP to position the phosphate groups for phosphoryl transfer (Figure 3C and Figure 3—figure supplement 1 and 2). Surprisingly, a pyrophosphate (PP_i_) molecule and two Ca^2+^ ions are bound within the kinase catalytic cleft (Figure 3D and E). PP_i_ is likely generated from ATP that was added to the crystallization condition. The presence of a PP_i_ molecule in the catalytic cleft indicates that SidJ may have an ATP-dependent catalytic function but not traditional phosphoryl transfer catalyzed by protein kinases.

**Figure 3.**
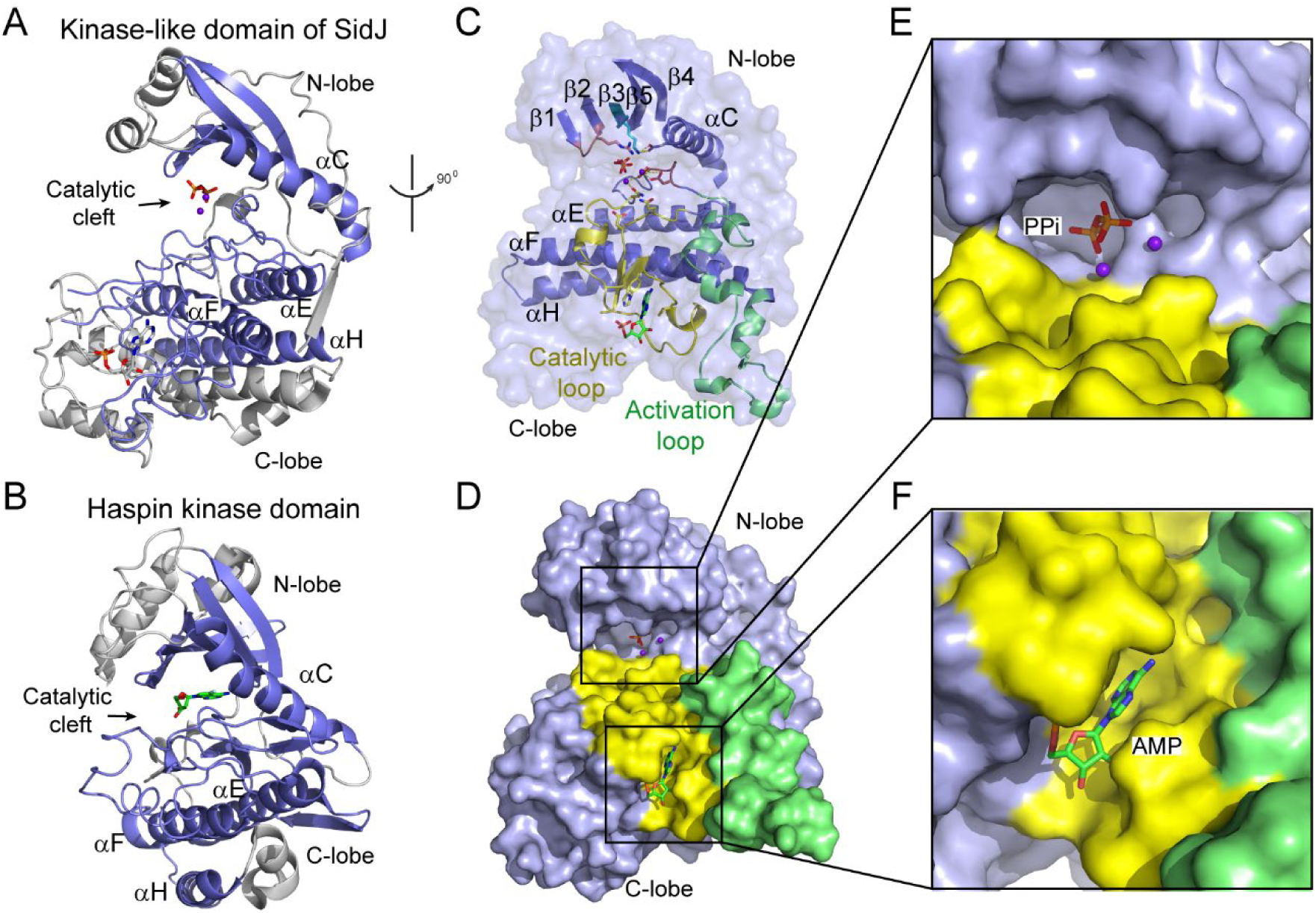
The core of SidJ adopts a protein kinase fold. (A) Cartoon diagram of the kinase-like domain of SidJ. Secondary structure elements that are conserved in protein kinases are colored in blue. Ca^2+^ ions are shown as purple spheres while the pyrophosphate and AMP molecules are shown in sticks. (B) Cartoon representation of the kinase domain of Haspin kinase (PDB ID: 2WB6). The conserved structural core colored in blue is displayed with a similar orientation to (A). (C) An orthogonal view of the conserved secondary structural elements in the SidJ kinase-like domain. The N-lobe is comprised of five antiparallel β-strands and αC helix. The C-lobe is primarily α helical. Secondary structural features are named according to PKA nomenclature. The activation loop is colored in green, the catalytic loop in yellow, and the glycine rich loop in salmon. Conserved residues within the kinase-like catalytic cleft are shown in sticks. (D) Surface representation of the SidJ kinase-like domain, depicting the catalytic cleft formed between the N- and C-lobes and the migrated nucleotide binding site formed mainly by residues within the catalytic loop (yellow). The activation loop (green) makes close contact with the catalytic loop. (E) Zoom-in view of catalytic clefts outlined in (D). The kinase catalytic cleft contains two Ca^2+^ ions and a pyrophosphate (PP_i_) molecule. (F) Expanded view of migrated nucleotide binding pocket bound with an AMP.

In contrast to the N-lobe, the C-lobe of the kinase-like domain is mainly helical. Three recognizable helices equivalent to the αE, αF, and αH helices in protein kinases set a foundation for three catalytic signature motifs on the C-lobe, including the HRD motif-containing catalytic loop, the DFG motif-containing Mg^2+^-binding loop, and the activation loop. These motifs are distributed within a long peptide connecting the αE and αF helices and are positioned at a similar location as in protein kinases (Figure 3C). Despite many conserved features between SidJ and canonical protein kinases, there are two unique features in the catalytic loop of SidJ. First, the aspartic acid in the HRD motif conserved in canonical kinases is notably different in SidJ, in which Q486 takes the position of D166 in PKA for the activation of substrates. Second, the catalytic loop of SidJ contains a 48-residue insertion between Q486 and the downstream conserved N534, albeit there are only four residues between D166 and N171 in PKA (Figure 3C and Figure 3—figure supplement 1 and 2). Interestingly, this large insertion creates a pocket that accommodates an AMP molecule (likely the breakdown product from ATP) (Figure 3D and F). The AMP molecule was also observed in this so-called migrated nucleotide-binding pocket in a recent reported SidJ-CaM structure (Black et al., 2019). The presence of this unique migrated nucleotide-binding pocket in SidJ further indicates that SidJ may have a distinct catalytic function other than a canonical protein kinase. Indeed, we were unable to detect any kinase activity for SidJ by in vitro kinase assays using [γ-^32^P]ATP (Figure 3—figure supplement 3A and 3B), even though most of the catalytic and scaffolding motifs essential for protein kinases are conserved in the SidJ kinase-like domain. In light of a recent discovery that the SelO pseudokinase has AMPylation activity (Sreelatha et al., 2018), we then tested whether SidJ is an AMPylase. A similar assay was performed with the substitution of ATP by [α-^32^P]ATP. Surprisingly, ^32^P incorporation was observed for SidJ itself but not for SdeA (Figure 3—figure supplement 3C and 3D). Interestingly, similar auto-AMPylation activity of SidJ was also observed in a recent publication (Gan et al., 2019). It is likely that auto-AMPylation of SidJ may be either a side reaction or an intermediate step for SidJ-mediated modification on SdeA.

### SidJ Catalyzes Polyglutamylation of SdeA

To determine the exact catalytic function of SidJ, we used a SILAC (Stable Isotope Labeling by Amino acids in Cell culture) mass spectrometry. HEK293T cells grown in complete medium containing heavy [^13^C6]lysine [^13^C6]arginine were co-transfected with GFP-SdeA and mCherry-SidJ while cells grown in regular medium were transfected with GFP-SdeA and a mCherry plasmid control. GFP-SdeA proteins were enriched by immunoprecipitation. MS analysis of immunoprecipitated SdeA revealed that one trypsinized peptide corresponding to the SdeA mono-ADP ribosylation catalytic site (residues 855-877) was dramatically reduced in the heavy sample prepared from cells transfected with both SidJ and SdeA compared to its light counterpart prepared from cells transfected with SdeA and a control plasmid (Figure 4A and B). This peptide generates two signature ions upon MS2 fragmentation due to the presence of two labile proline residues in the sequence. We then used this feature to search for any peptide from the heavy sample that produced these two signature ions. Multiple MS2 spectrum contained these two signature ions (Figure 4—figure supplement 1A-D). Strikingly, all these peptides had a mass increase of *n* x 129 Da, which matches the mass change corresponding to posttranslational modification by polyglutamylation. The MS data were then re-analyzed for polyglutamylation. The modification of the SdeA peptide was revealed as either mono-, di-, or tri-glutamylation with the predominant species being di-glutamylation (Figure 4—figure supplement 1E). Furthermore, the polyglutamylation site was identified at SdeA residue E860 by MS/MS analysis (Figure 4C). The activity of SidJ was then reconstituted in vitro using [U-^14^C]Glu. Consistent with the mass spectrometry results, wild type SdeA core (residues 211-1152) but not its E860A mutant, was modified with glutamate. In addition, polyglutamylation of SdeA by SidJ was dependent on both CaM and ATP/Mg^2+^ (Figure 4D). Since E860 is one of the key catalytic residues in the mART domain of SdeA (Figure 4—figure supplement 2), it is likely that polyglutamylation of E860 may inhibit SdeA-mediated ADP-ribosylation of ubiquitin.

**Figure 4.**
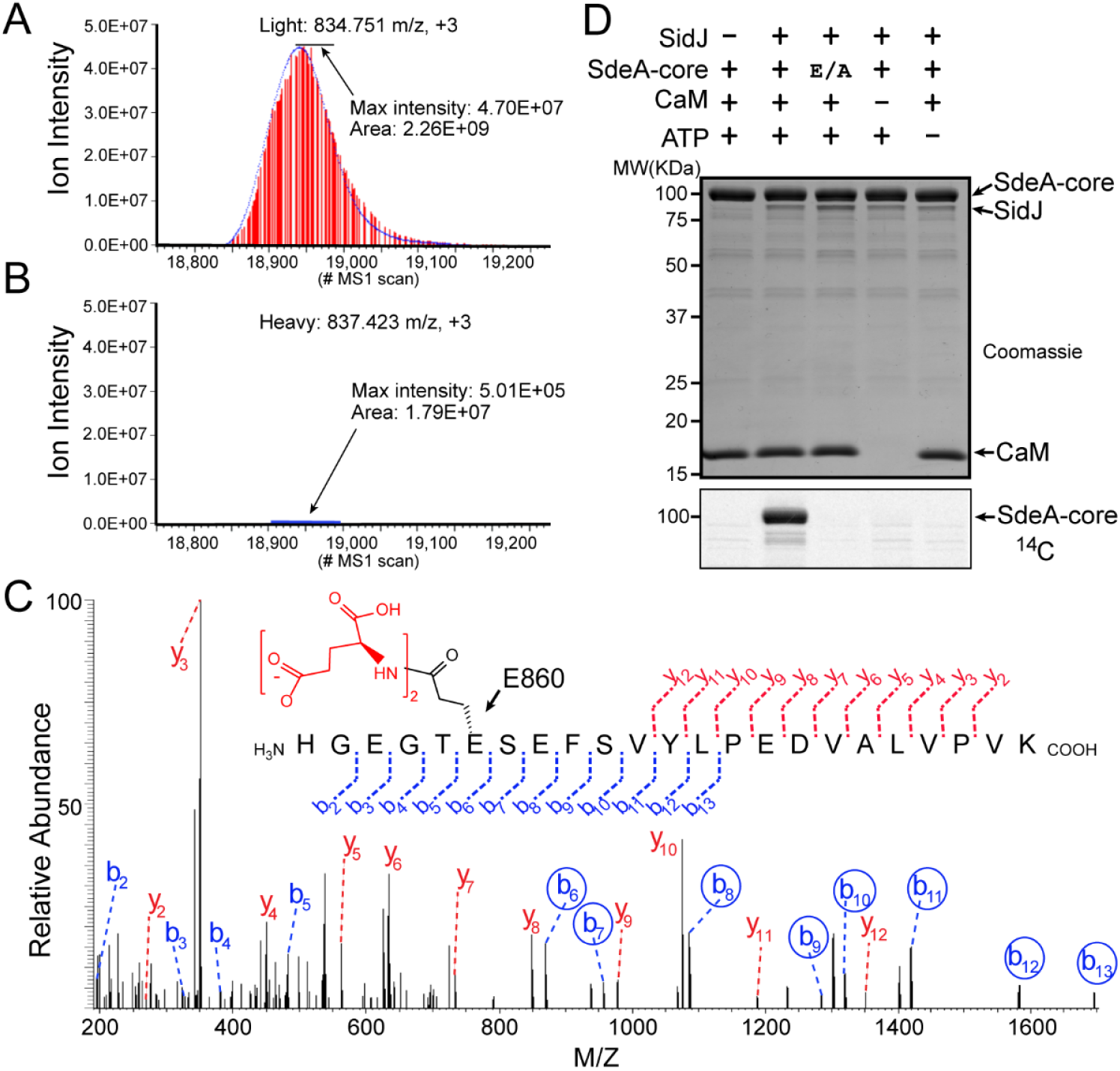
SidJ catalyzes polyglutamylation of SdeA. Reconstructed ion chromatograms for the SdeA peptide (residues 855-877) from (A) cells grown in light medium and co-transfected with GFP-SdeA and mCherry vector control or (B) cells grown in heavy medium and co-transfected with GFP-SdeA and mCherry-SidJ. (C) MS2 spectrum of a di-glutamylated SdeA mART peptide with b ions labeled in blue and y ions labeled in red. The peptide sequence corresponding to fragmentation is depicted above the spectrum. Circled B ions represent a mass increase corresponding to diglutamylation (258.085 Da). (D) In vitro glutamylation of SdeA with [U-^14^C]Glutamate. E/A corresponds to the E860A point mutant of SdeA. Proteins were separated by SDS-PAGE and visualized with Coomassie stain (top panel) and autoradiogram of the gel is shown in bottom panel.

### SidJ Suppresses the PR-ubiquitination Activity of SdeA

To test whether SidJ directly inhibits SdeA activity, we performed in vitro ubiquitin modification and PR-ubiquitination assays with either untreated or SidJ-pretreated SdeA. Ubiquitin was modified in the presence of NAD^+^ by purified SdeA to generate ADPR-Ub as indicated by a band-shift of ubiquitin on a Native PAGE gel. However, when SdeA was pre-incubated with SidJ, CaM, ATP/Mg^2+^, and glutamate, ubiquitin modification by SdeA was substantially reduced (Figure 5A) but was not affected if the pretreatment lacked either glutamate, ATP or CaM (Figure 5A). In agreement with impaired ADPR-Ub generation, SdeA-mediated PR-ubiquitination of a substrate, Rab33b was also inhibited in a reaction with SidJ-treated SdeA (Figure 5B). We further investigated whether SidJ can also regulate the PR-ubiquitination process during *Legionella* infection. HEK293T cells were first transfected with 4xFlag-tagged Rab33b and FCγRII then infected with the indicated opsonized *Legionella* strains for 2 hours. Rab33b was then immunoprecipitated and analyzed with anti-Flag Western blot (Figure 5C). The total amount of PR-ubiquitinated Rab33b was more than doubled in cells infected with *ΔsidJ* strain. However, complementation with a plasmid expressing wild type, but not the D542A SidJ mutant, reduced Rab33b PR-ubiquitination to a level comparable to infection with the wild type *Legionella* strain (Figure 5C and D). Taken together, these data suggest that SidJ suppresses the PR-ubiquitination via SidJ-mediated polyglutamylation of SdeA.

**Figure 5.**
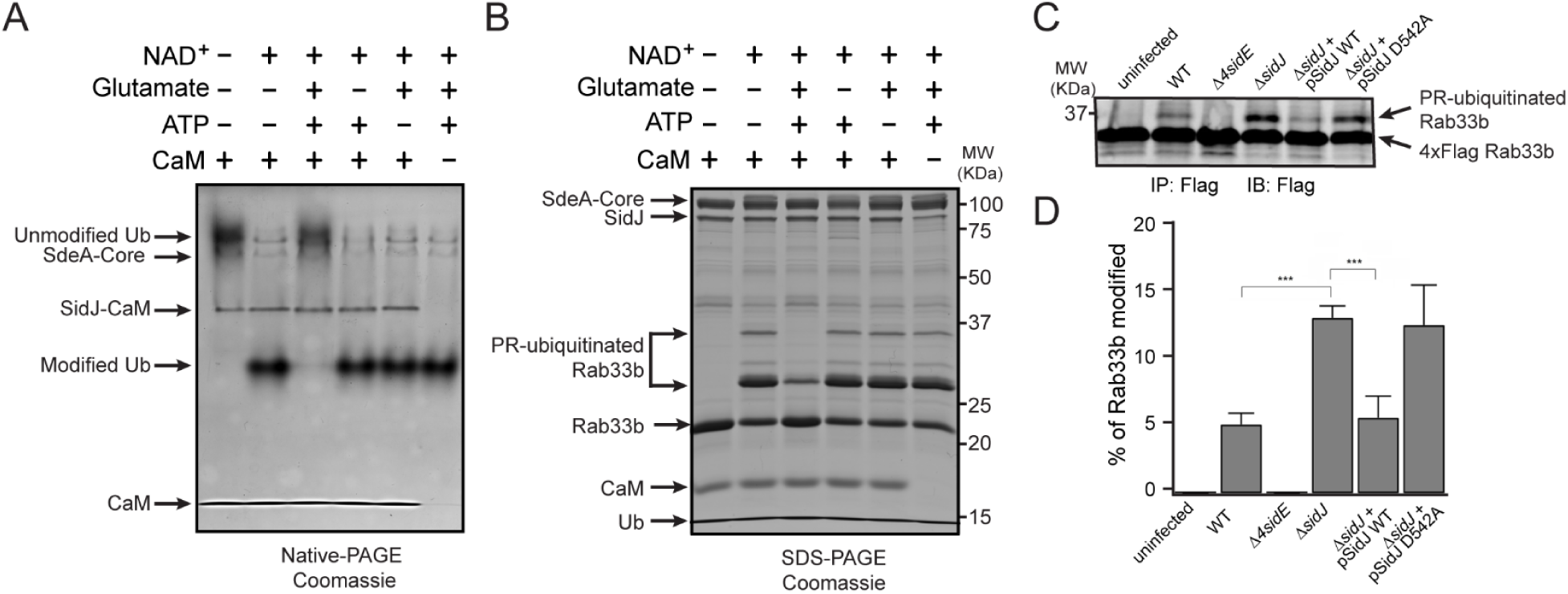
SidJ suppresses the PR-ubiquitination activity of SdeA. (A) SdeA Core was first incubated with SidJ for 30 minutes at 37°C with MgCl_2_, ATP, CaM, and in the presence or absence of glutamate. Then the SdeA mediated ADP-ribosylation of Ub was initiated by addition of Ub and NAD^+^ to the reaction mixture and further incubated for 30 minutes at 37°C. Final products were analyzed by Native PAGE to monitor the modification of Ub as an indirect readout for the polyglutamylation activity of SidJ. (B) In vitro SdeA PR-ubiquitination of Rab33b after a similar pretreatment by SidJ as in (A). The final products were analyzed by SDS-PAGE to monitor the generation of PR-ubiquitinated Rab33b. (C) PR-ubiquitination of Rab33b was increased in cells infected with Δ*sidJ L. pneumophila* strain. HEK293T cells expressing FCγRII and 4xFlag-Rab33b were infected with indicated *L. pneumophila* strains for 2 hours. 4xFlag-Rab33b proteins were enriched by anti-Flag immunoprecipitation and analyzed by anti-Flag Western blot. (D) Quantification of percentage of PR-ubiquitinated Rab33b in blots in panel C. Data are shown as means ± SEM of three independent experiments. ***P<0.001.

### Molecular Determinants of Protein Glutamylation Catalyzed by SidJ

The identification of SidJ as a polyglutamylase raised an intriguing question, how can a kinase-like enzyme attach glutamates to its targets. To address this question, selected residues in the canonical kinase catalytic cleft and in the migrated nucleotide binding pocket were mutagenized and the functions of these mutants were interrogated for their polyglutamylation activities and their ability to inhibit SdeA in vitro. In the SidJ kinase catalytic cleft, two Ca^2+^ ions are coordinated by residues N534, D542, and D545, while the PP_i_ molecule is stabilized by R352 from the Gly-rich loop and the conserved K367, which in turn is stabilized by E381 from the αC helix (Figure 6A and B). On the other hand, in the migrated nucleotide binding pocket, the aromatic adenine base of AMP is stacked with the imidazole ring of H492, while Y506 forms the interior wall of the pocket (Figure 6C). These residues were mutated to Alanine and the polyglutamylation activity of these SidJ mutants were examined. The polyglutamylation activity of SidJ was completely abolished in the K367A, D542A, and H492 mutants and was severely impaired in the N534A mutant. The polyglutamylation activity was slightly reduced in the R352A and Y506A mutants while the E381A, D489A, and D545A mutations had little or no impact on the activity of SidJ (Figure 6D-F). In addition, the polyglutamylation activity of SidJ mutants correlated well with their inhibition on SdeA-mediated modification of Ub (Figure 6—figure supplement 1).

**Figure 6.**
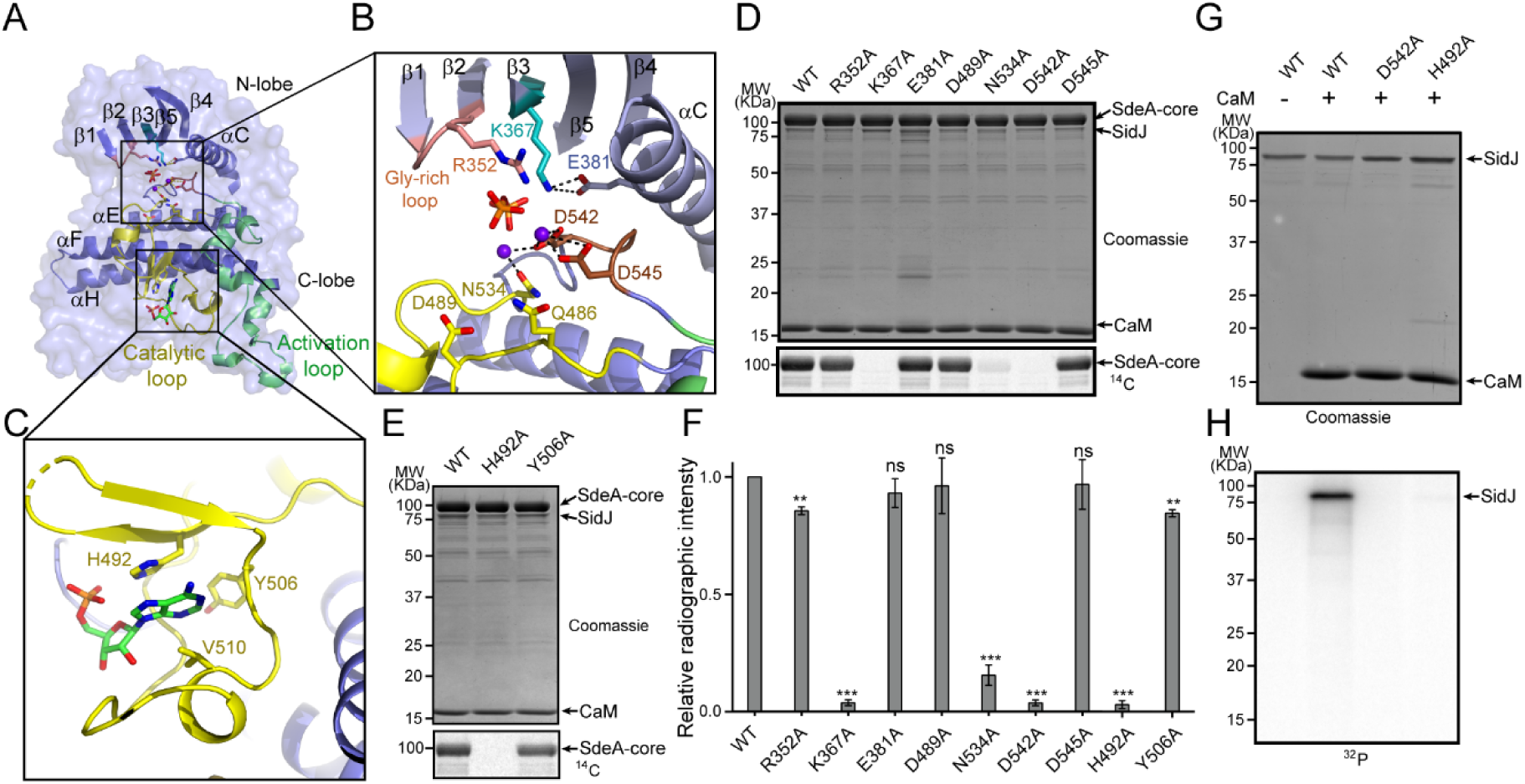
Molecular determinants of SidJ-mediated polyglutamylation. (A) Overall structure of SidJ kinase-like domain. (B) Enlarged view of kinase catalytic site of SidJ. Key catalytic residues are displayed in sticks. Pyrophosphate is shown as sticks and two calcium ions are shown as purple spheres. (C) Enlarged view of “migrated” nucleotide binding site with AMP displayed as sticks. (D) In vitro glutamylation of SdeA by SidJ active site mutants with [U-^14^C]glutamate after 15 minute reaction at 37°C. The proteins in the reactions were visualized by SDS-PAGE followed by Coomassie staining (top panel) and the modification of SdeA was detected by autoradiography (bottom panel). (E) In vitro glutamylation of SdeA by SidJ nucleotide binding site mutants with [U-^14^C]glutamate. The proteins in the reactions were analyzed by SDS-PAGE (top) and the glutamylation of SdeA was detected by autoradiography (bottom). (F) Quantification of the relative autoradiographic intensity of modified SdeA. Data are shown as means ± STD of three independent experiments. ns: not significant; **P<0.01; ***P<0.001. (G) SidJ and SidJ mutants were incubated with [α-^32^P]ATP and MgCl_2_ in the presence or absence of CaM. Representative SDS-PAGE gel was stained with Coomassie. (H) Autoradiogram of the gel in G to show the auto-AMPylation of SidJ.

It is intriguing that the polyglutamylation activity of SidJ was abolished by mutations at either the canonical kinase-like active site (K367A or D542A) or at the migrated nucleotide binding site (H492). It has been proposed that the kinase-like active site catalyzes the transfer of AMP from ATP to E860 on SdeA while the migrated nucleotide binding site catalyzes the replacement of AMP with a glutamate molecule to complete glutamylation of SdeA at E860 (Black et al., 2019). However, it may also be possible that the glutamylation reaction takes place at the kinase-like active site whereas the migrated nucleotide binding site serves as an allosteric site, in which binding of an AMP molecule at the migrated nucleotide binding site is a prerequisite for SidJ activation. To test these two hypothesis, we took advantage of the auto-AMPylation activity of SidJ. If the first hypothesis is true, one would expect that the SidJ H492A mutant would be competent for auto-AMPylation since it has an intact kinase active site. Strikingly however, SidJ auto-AMPylation was completely abolished in both the D542A and H492A mutants. These data suggest that the migrated nucleotide binding site is likely an allosteric site (created entirely by a large insertion within the catalytic loop). The binding of a nucleotide to this site is likely to stabilize the catalytic loop of the kinase-like domain in a catalytically competent conformation.

### Activation of SidJ by CaM

Our in vitro assays demonstrated that the polyglutamylation activity of SidJ requires binding with CaM. We next asked how CaM activates SidJ. A close examination of the SidJ-CaM complex structure revealed that the highly acidic CaM binds with the basic IQ-helix of SidJ mainly through its C-lobe (Figure 7A and C and Figure 7—figure supplement 1). The C-lobe of CaM assumes a semi-open conformation, which creates a groove between CaM helices F and G and helices E and H to grip the amphipathic IQ-helix of SidJ (Figure 7—figure supplement 2). Conserved hydrophobic residues aligned inside the groove make numerous van der Waals interactions with the hydrophobic side of the IQ-helix centered at I841, whereas acidic residues located at the edge of the groove form hydrogen bonds and salt bridges with polar residues on the hydrophilic side of the IQ-helix (Figure 7C). In contrast, the N-lobe of CaM maintains a closed conformation similar to that observed in free apo-CaM (Kuboniwa et al., 1995) or in the myosin V IQ1-CaM complex (Houdusse et al., 2006) even though one Ca^2+^ ion is chelated by the first EF-hand of CaM (Figure 7—figure supplement 3A). Interestingly, the binding of this calcium ion does not cause a conformational change observed in CaM fully chelated with Ca^2+^ (Meador et al., 1992) since the conserved E31 of CaM is not positioned for chelation at the -Z coordination position (Figure 7—figure supplement 3B-D).

**Figure 7.**
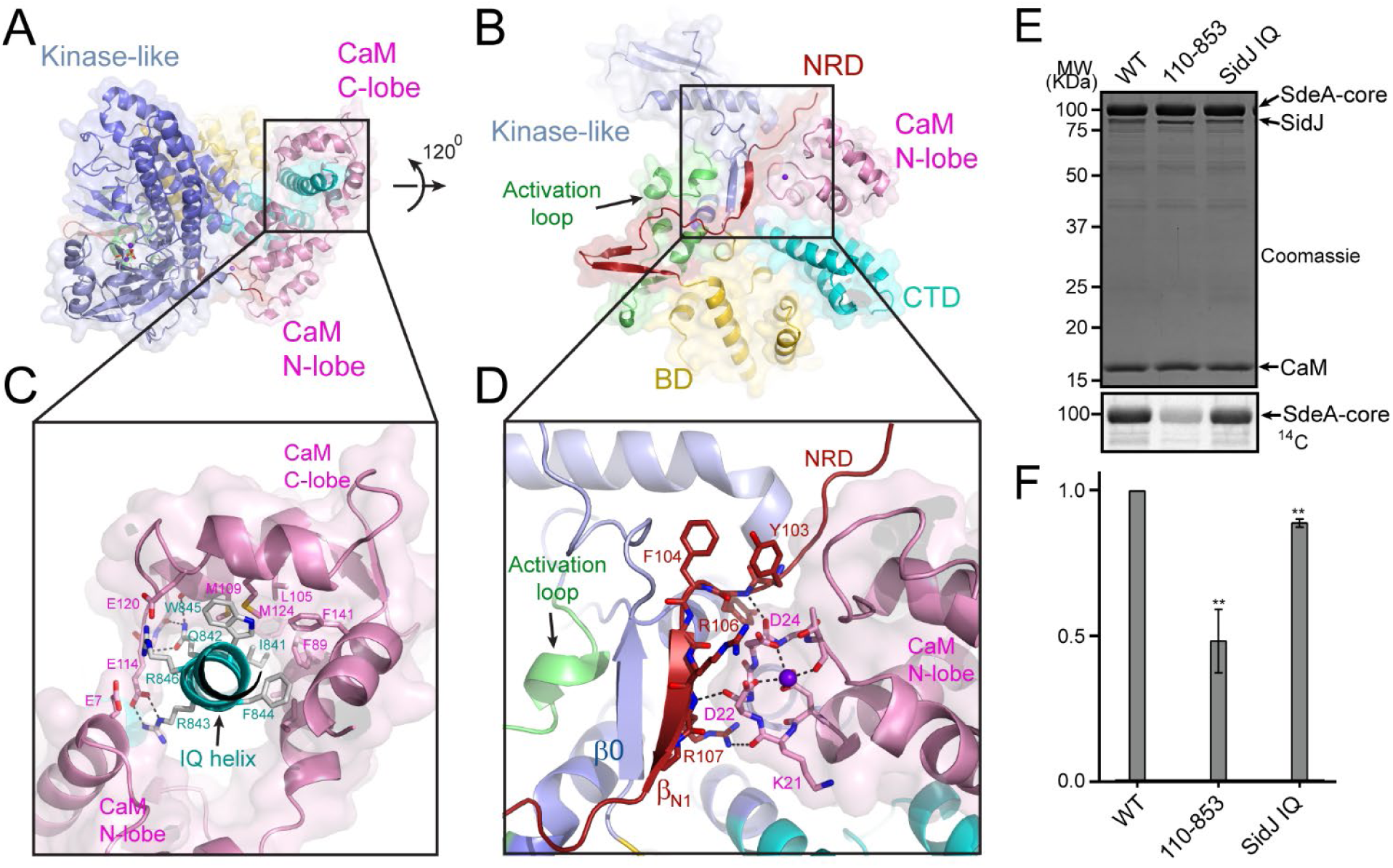
Activation of SidJ by CaM. (A) The structure of the SidJ-CaM complex showing the C-lobe of CaM (pink) “gripping” the IQ-motif helix (cyan) of SidJ. (B) A 120° rotated view of the complex in panel A showing the N-lobe of CaM contacts with the NRD domain (maroon) of SidJ (C) Enlarged view of interface between the SidJ IQ helix and the C-lobe of CaM. Residues involved in the interactions between the IQ helix and CaM are depicted as sticks. Hydrogen bonds and electrostatic interactions depicted with dashed lines. (D) Enlarged region of interface between the NRD and CaM. Purple sphere represents the Ca^2+^ ion bound to CaM. (E) In vitro glutamylation of SdeA by SidJ mutants. The proteins in the reactions were visualized by SDS-PAGE followed by Coomassie staining (top panel) and the modification of SdeA was detected by autoradiography (bottom panel). (F) Quantification of the relative autoradiographic intensity of modified SdeA. Data are shown as means ± STD of three independent experiments. **P<0.001.

A structural comparison of SidJ-CaM with myosin V IQ1-CaM complex revealed that although both of the N- and C-lobes of CaM have a similar conformation with their corresponding lobes, the relative orientation between two lobes assumes a remarkably different conformation in the two complexes (Figure 7—figure supplement 4). Unlike the CaM in the myosin V IQ1-CaM complex, in which the N-lobe packs close to and makes a large number of contacts with the IQ-helix, the CaM N-lobe in the SidJ-CaM complex is shifted away from the IQ-helix and engages extensive interactions with other basic areas of SidJ, including the first β-strand (β_N1_) of the NRD domain (Figure 7B and D). Besides electrostatic interactions between the CaM N-lobe and SidJ, two carbonyl groups within the first Ca^2+^-binding loop of the CaM N-lobe form hydrogen bonds with two backbone amide groups of the β_N1_ strand (Figure 7D). These interactions between the CaM N-lobe and the β_N1_ strand may stabilize a two-stranded antiparallel β sheet composed of β_N1_ of the NRD domain and β0 of the kinase-like domain, which may further stabilize the activation loop in a presumably active conformation (Figure 7D). The stabilization of the activation loop is reminiscent of the activation process in canonic kinases, in which phosphorylation of specific residues in the activation loop provides an anchor to maintain the activation loop in the correct conformation for catalysis (Adams, 2003). Based on these structural observations, we hypothesized that CaM-binding stabilizes a two-stranded β sheet on the surface of SidJ, which in turn interacts with the activation loop of the kinase-like domain to maintain the activation loop in an active conformation. Indeed, although the SidJ IQ mutant demonstrated a modest reduction in activity, the β_N1_ deleted SidJ truncation (residue 110-853) showed a severe impairment in polyglutamylation of SdeA (Figure 7E and F). Together, our data suggest that CaM-binding may activate SidJ through a network of interactions involving the CaM N-lobe, the β_N1_ strand of the NRD, and the β0 strand and the activation loop of the kinase-like domain.

## Discussion

In this study, we reported the crystal structure of a *Legionella* effector SidJ in complex with human CaM. Through structural, biochemical, and mass spectrometric studies, we identified the biochemical function of SidJ as a protein polyglutamylase that specifically adds glutamates to a catalytic glutamate residue E860 of another *Legionella* effector SdeA and thus inhibits the PR-ubiquitination process mediated by SdeA. To date, the only enzymes that have been identified to catalyze protein glutamylation belong to the tubulin tyrosine ligase-like (TTLL) protein family (Janke et al., 2008). The TTLL enzymes have an active site that lies between two characteristic α/β domains. An elegant crystal structure of TTLL7 in combination with a cryo-electron microscopy structure of TTLL7 bound to the microtubule revealed that the anionic N-terminal tail of β-tubulin extends through a groove towards the ATP-binding active site for the modification. By contrast, the catalytic core of SidJ adopts a protein kinase-like fold. Surprisingly, besides the canonical kinase-like active site, SidJ also has a second “migrated” nucleotide-binding site created by a large insertion in the kinase catalytic loop. The two sites are both required to complete the polyglutamylation reaction as single amino acid point mutations of key residues at either site inactivate SidJ (Figure 6). We further showed that the auto-AMPylation activity of SidJ was also impaired by mutations at either the kinase-like active site or the migrated nucleotide-binding site. These observations led us to propose a reaction model for SidJ-mediated polyglutamylation (Figure 8). In this model, SidJ is activated by CaM binding at its C-terminal IQ helix and a nucleotide binding at its migrated nucleotide-binding pocket. Activated SidJ first attaches the AMP moiety from ATP to the γ-carbonyl group of residue E860 of SdeA. In the second step, the adenylated E860 is attacked by the amino group of a free glutamate to form an isopeptide linkage by releasing AMP. However, several prominent questions remain to be addressed, such as how SidJ recognizes SdeA and specifically attaches glutamates to residue E860 of SdeA. Furthermore, how the specificity is achieved to modify SdeA with glutamate residues but not other amino acids. To answer these questions, more biochemical assays, as well as structural studies of SidJ in complex with substrates or intermediates are warranted.

**Figure 8.**
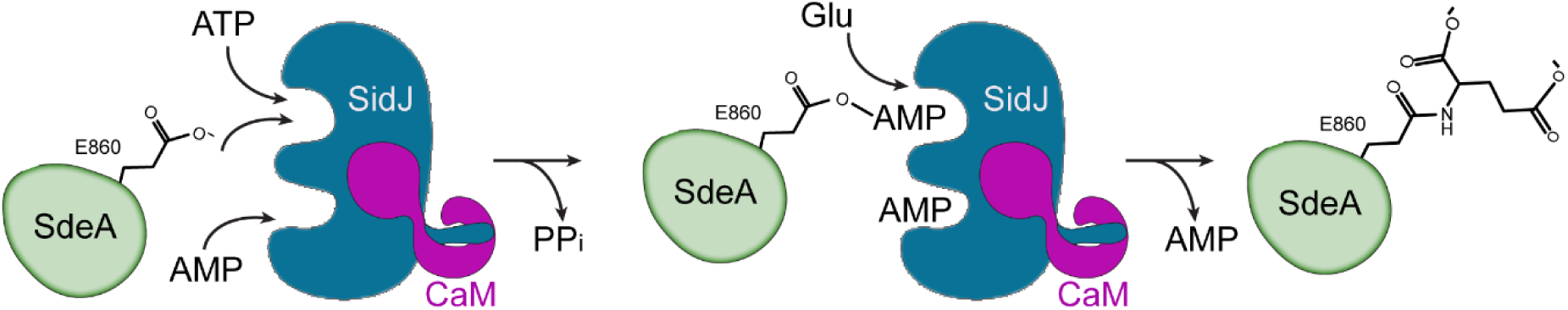
Hypothetic reaction model for SidJ-mediated polyglutamylation of SdeA. SidJ has a kinase-like catalytic cleft, a regulatory nucleotide-binding pocket and C-terminal CaM-binding IQ helix. Binding of a nucleotide to the allosteric regulatory site and CaM with the IQ motif activates SidJ. SidJ mediated SdeA polyglutamylation involves two steps. In the first step, SidJ AMPylates SdeA by transferring the AMP moiety from ATP to the γ-carbonyl group of SdeA E860 and releasing a pyrophosphate molecule. In the second step, a glutamate molecule is activated at the kinase active site and its amino group serves as a nucleophile to attack the AMPylated E860. As a result, this glutamate is conjugated to E860 through an isopeptide bond and an AMP molecule is released.

Interestingly, SidJ contains a C-terminal consensus IQ motif that mediates CaM-binding independent of calcium. The binding of the IQ helix is mainly through the CaM C-lobe, which adopts a semi-open conformation similar to that observed in apo-CaM-IQ helix complex. However, unlike other apo-CaM complexes, where the CaM N-lobe wraps around the IQ-helix, the N-lobe in the SidJ-CaM complex rotates along the inter-lobe linker about 120 degree and swings away from the IQ-helix to engage in extensive interactions with other parts of SidJ, particularly the β_N1_ strand of the NRD domain. These interactions may allosterically stabilize a hydrophobic core, which may serve as an anchor point for the kinase activation loop to active the enzyme. SidJ-CaM seems to apply a unique CaM-dependent regulatory mechanism to maintain an active conformation. Thus, the binding mode of CaM with SidJ presents an exemplary mechanism to the repertoire of CaM-effector interactions. The activation of SidJ by CaM is also of particular interest from an evolutionary point of view. Both SidJ and SdeA are expressed in *Legionella* cells, while the polyglutamylation, hence the inhibition of SdeA can only occur after they have been delivered into eukaryotic host cells. A similar example has been reported for the CaM-mediated activation of anthrax adenylyl cyclase exotoxin (Drum et al., 2002). This type of cross-species regulation may represent a common theme in bacterial pathogen-eukaryotic host interactions.

SidJ was first identified as a metaeffector that neutralizes the toxicity of the SidE family phosphoribosyl ubiquitin ligases in yeast (Havey and Roy, 2015; Jeong et al., 2015). A previous publication assigned SidJ as a deubiquitinase that deconjugates phosphoribosyl-linked protein ubiquitination (Qiu et al., 2017). However, this unusual deubiquitination activity was not repeatable in another study (Black et al., 2019), as well as in our unpublished studies. The definitive biochemical function of SidJ is now revealed in this study, as well as by recent reports (Bhogaraju et al., 2019; Black et al., 2019; Gan et al., 2019), as a polyglutamylase that adds glutamates to a specific catalytic residue E860 of SdeA and subsequently inhibits the PR-ubiquitination activity of SdeA. An interesting question arises at this point as whether there are other glutamylation substrates, especially from the host, besides the SidE family PR-ubiquitination ligases. Given that SidJ is one of the few effectors that exhibits growth defects when deleted from *L. pneumophila* and plays a role in membrane remodeling during *Legionella* infection (Liu and Luo, 2007), it is possible that SidJ modifies host targets to control certain host cellular processes. On the other hand, it seems to be a common scheme in *Legionella* species to encode effectors catalyze counteractive reactions. For example, the *Legionella* effector SidM/DrrA AMPylates Rab1 and locks it in an active GTP state (Muller et al., 2010) while SidD is a deAMPylase that antagonizes SidM (Neunuebel et al., 2011; Tan and Luo, 2011). Another example is the pair of effectors AnkX and Lem3, of which AnkX transfers a phosphocholine moiety to Rab1 family members (Mukherjee et al., 2011) while Lem3 removes the phosphocholine moiety added by AnkX from Rab1 (Tan et al., 2011). In respect of this scheme, it is possible that *L. pneumophila* may also encode an effector that counteracts SidJ by removing glutamate residues from targets. Future investigation of effectors harboring such a de-glutamylation activity would provide a comprehensive understanding of the regulation cycle of protein glutamylation taken place during *Legionella* infection.

It is also noteworthy that homologs of SidJ can be detected in a variety of microorganisms, including *Elusimicrobia bacterium, Desulfovibrio hydrothermalis, and Waddlia chondrophila*. Furthermore, the key catalytic motifs found in SidJ are also readily detectable in these homologs (Figure 3—figure supplement 1). It would be interesting to elucidate whether these SidJ homologs have a comparable activity to SidJ. In summary, our results have identified SidJ contains kinase-like fold and functions as a protein polyglutamylase. Our results may contribute inspiring hints to the search for other potential protein polyglutamylases and to the studies of pseudokinases for alternative ATP-dependent activities.

## Materials and Methods

### Cloning and mutagenesis

SidJ was PCR amplified from *Legionella* genomic DNA and digested with BamH1 and Sal1 and cloned into pmCherry-C1 and pZL507 (obtained from Dr. Zhao-Qing Luo, Purdue University) vectors for mammalian and *Legionella* expression respectively. For protein purification, the SidJ 89-853 truncation was amplified for pmCherry and cloned into the vector pET28a-6xHisSumo using vector BamH1 and the reverse isoschizomer Xho1 site. Human CaM2, SdeA 211-1152 truncation was cloned into pET28a 6xHisSumo using BamH1 and Xho1 sites on both vector and insert. For *Legionella* genomic deletions, 1.2 Kb regions upstream and downstream of SidJ were cloned into the pRS47s suicide plasmid (obtained from Dr. Zhao-Qing Luo). Site directed mutagenesis was then performed with overlapping primers on each vector. All constructs were transformed into chemically competent Top10 cells, with the exception of pRS47s vector which was transformed into DH5α λpir.

### Protein Purification

All pET28a 6x-HisSumo constructs including SidJ 89-853, CaM, SdeA 211-1152, and point mutants were transformed into E. coli Rosetta (DE3) cells. Single colonies were then cultured in Luria-Bertani (LB) medium containing 50 µg/ml kanamycin to a density between 0.6 and 0.8 OD_600_. Cultures were induced with 0.1 mM isopropyl-B-D-thiogalactopyranoside (IPTG) at 18°C for 12 hours. Cells were collected by centrifugation at 3,500 rpm for 15 minutes at 4°C and sonicated to lyse bacteria. To separate insoluble cellular debris, lysates were then centrifuged at 16,000 rpm for 45 minutes at 4°C. The supernatant was incubated with cobalt resin (Gold Biotechnology) for 2 hours at 4°C to bind proteins and washed extensively with purification buffer (20 mM Tris pH 7.5, 150 mM NaCl). Proteins of interest were then digested on the resin with SUMO-specific protease Ulp1 to release the protein from the His-SUMO tag and resin. The digested protein was concentrated and purified further by FPLC size exclusion chromatography using a Superdex S200 column (GE life science) in purification buffer. Pure fractions were collected and concentrated in Amicon Pro Purification system concentrators.

### Native PAGE analysis of SidJ-CaM complex

SidJ 89-853 WT and SidJ IQ mutant were incubated at a concentration of 5 μM with 10 μM of CaM in the presence of 1 mM CaCl_2_ or 1 mM EGTA in 50 mM Tris pH 7.5 and 150 mM NaCl. Samples were then analyzed by Native PAGE and gels were stained with Coomassie Brilliant Blue.

### Isothermal titration calorimetry (ITC)

SidJ 89-853 WT, IQ mutant and CaM were used for ITC experiments. CaM at 88.6 µM concentration was titrated into SidJ 89-853 WT and IQ mutant at 20 µM concentration. CaM was titrated in 15 injections at 5 µL with spacing between injections ranging from 150 s to 400 s, until the baseline equilibrated. These experiments used the Affinity ITC from TA instruments at 25°C. Data analysis was performed on NanoAnalyze v3.10.0.

### Analytical size exclusion

SidJ and IQ mutant were incubated at a concentration of 35 μM in the presence or absence of 1.2 molar ratio of CaM. 125 μL of solution were injected onto a Superdex 200 Increase 100/300 GL column (GE) and separated at 0.7 mL/min on an AKTA Pure 25L System (GE). UV traces were generated using R-Studio Software and 0.5 mL fractions were collected and analyzed by SDS-PAGE. Gels were stained with Coomassie brilliant blue.

### Protein crystallization

Protein crystallization screens were performed on the Crystal Phoenix liquid handling robot (Art Robbins Instruments) at room temperature using commercially available crystal screening kits. Prior to screening and hanging drop experiments, SidJ and CaM were incubated at a 1 to 2 molar ratio for 1 hour on ice. The conditions that yielded crystals from the screen were optimized by hanging-drop vapor diffusion by mixing 1 µL of the protein complex with 1 µL of reservoir solution. All optimization by hanging-drop vapor diffusion was performed at room temperature. Specifically, for SidJ-CaM crystallization, SidJ was concentrated to 9.4 mg/mL and crystallized in 0.2 M sodium iodide, 15% PEG 3350, 0.1 M Tris pH 9.2, 1 mM CaCl_2_ and 1 mM ATP. Rod shaped crystals formed within 4-5 days.

### X-Ray diffraction data collection and processing

Diffraction datasets for SidJ-CaM were collected at National Synchrotron Light Source II (NSLSII) beamline AMX (17-ID-1) at Brookhaven National Laboratory. Before data collection, all crystals were soaked in cryoprotectant solutions that contained the crystallization reservoir condition, supplemented with 25% glycerol. All soaked crystals were flash frozen in liquid nitrogen prior to data collection. X-ray diffraction data were indexed, integrated, and scaled with HKL-2000 (Otwinowski and Minor, 1997).

### Structure determination and refinement

The structure of SidJ was solved by using single wavelength anomalous dispersion (SAD) method with selenomethionine-incorporated crystals. Heavy atom sites were determined and phasing was calculated using HKL2MAP (Pape and Schneider, 2004). Iterative cycles of model building and refinement were performed using COOT (Emsley and Cowtan, 2004) and refmac5 (Murshudov et al., 1997) of the CCP4 suite (Collaborative Computational Project, 1994). Surface electrostatic potential was calculated with the APBS (Baker et al., 2001) plugin in PyMOL. All structural figures were generated using PyMOL (The PyMOL Molecular Graphics System, Version 1.8.X, Schrödinger, LLC).

### Protein sequence analysis

Sequences homologous to SidJ were selected from the NCBI BLAST server. All sequences were aligned using Clustal omega (Sievers et al., 2011) and colored using the Multiple Align Show server (http://www.bioinformatics.org/sms/index.html)

### SILAC and mass spectrometry sample preparation

HEK293T cells were grown for 5 passages in media containing Light (^12^C^14^N Lys and Arg), or heavy (^13^C^15^N Lys and Arg) amino acids. Light HEK-293T cells transfected for 36 hours with pEGFP-SdeA and pmCherry and heavy HEK-293T cells transfected with pEGFP-SdeA and pmCherry-SidJ. Cells were then washed twice with cold PBS and resuspended using a cell scraper into lysis buffer (50 mM Tris pH 8.0, 150 mM NaCl, 1% Triton X-100, 0.1 % NaDOC, PMSF and Roche Protease Cocktail). Cells were sonicated and lysates were centrifuged at 16,000xg for 15 minutes at 4°C. Supernatants were incubated for 4 hours with GFP nanobeads and washed with IP wash buffer (50 mM Tris-HCl pH 8.0, 150 mM NaCl, 1% Triton). Proteins were eluted by incubation of resin in 100 mM Tris HCl pH 8.0, 1% SDS at 65°C for 15 minutes. Eluates were reduced with 10 mM DTT and alkylated with 25 mM iodoacetamide. Heavy and light samples were mixed and precipitated on ice in PPT (49.9% ethanol, 0.1% glacial acetic acid, and 50% acetone). Proteins were pelleted by centrifugation at 16,000xg, dried by evaporation and resolubilized in 8 M Urea in 50 μM Tris pH 8.0. The sample was digested overnight with trypsin gold at 37°C. Trypsinized samples were acidified with formic acid and triflouroacetic acid and bound to a C18 column (Waters) and washed with 0.1% acetic acid. Peptides were eluted with 80% acetonitrile and 0.1% acetic acid and dried. Samples were resuspended in 0.1 picomol/uL of angiotensin in 0.1% TFA and frozen for mass spectrometry analysis.

### Mass spectrometry analysis

Trypsinized SILAC-IP eluates from HEK-293T cells expressing either GFP-SdeA grown in ^12^C^14^N Lys + Arg, or GFP-SdeA and mCherry-SidJ grown in ^13^C^15^N Lys + Arg were analyzed on a ThermoFisher Q-Exactive HF mass spectrometer using a homemade C_18_ capillary column. Peptide spectral matches were identified using a SEQUEST search through Sorcerer2 from Sage-N, and subsequently quantified by Xpress to identify peptides that were highly enriched in the SdeA-light sample (indicating the absence of that peptide from the heavy condition because of a modification). Following identification of a single peptide, R.HGEGTESEFSVYLPEDVALVPVK.V, that was disproportionately enriched in the SdeA-only condition, the .raw file from the mass spectrometer was manually inspected to find MS2 spectra which had a similar retention time and contained peaks at m/z = 351 and 1074, as these masses were characteristic of the precursor peptide found in the SdeA-only condition due to the peptide containing two labile prolines. The monoisotopic precursor mass of the original, unmodified peptide from the SdeA-only condition was subtracted from the precursor mass of the most abundant peak fitting the above description. This difference corresponded to glutamylation. The original file was subsequently searched in Sorcerer2 using glutamylation (monoisotopic mass of 129.042587 Da) as a differential modification, and glutamylation sites were identified in the original peptide with Xpress scores that corresponded to their presence exclusively in the heavy condition (SdeA + SidJ).

### In vitro glutamylation assays and SdeA inhibition

In vitro glutamylation assays were conducted with 0.5 μM SidJ 89-853, 5 μM CaM, 5 mM MgCl_2_, 5 mM Glutamatic Acid, and 1 μM SdeA 231-1152 in a buffer containing 50 mM Tris pH 7.5 and 50 mM NaCl. Reactions were then initiated by addition of 1 mM ATP for 30 minutes at 37°C. For SdeA inhibition assays, a second ubiquitination reaction was conducted containing 25 μM ubiquitin and initiated with 1 mM NAD^+^. When testing PR-Ub ligation 10 mM Rab33b 1-200 served as a substrate. Reactions were then fixed with 5X SDS loading buffer or 6X DNA loading buffer and electrophroresed with 12% SDS-PAGE gels to assay PR-Ubiquitination, or native gels to assay Ub modification. Gels were stained with Coomassie Brilliant Blue stain.

### Radioactive glutamylation assay

Assays were conducted in a similar manner to non-radioactive glutamylation assays with the following exceptions, the concentration of SdeA was 2 μM, and 50 μM (1.76 nCi) of U-C14 Glutamate (Perkin Elmer) was used as a reactant. For SidJ mutants, glutamylation reactions were terminated after a 15 min reaction at 37°C with 5X SDS loading buffer. Samples were then electrophoresed by SDS-PAGE and gels were dried. Protein labeling was then visualized by a 3-4 days exposure using an image screen (FUJI BOS-III) and a phosphoimager (Typhoon FLA 7000, GE). Quantifications were performed using the program FIJI where background signal was subtracted from band intensity and divided by wild type SidJ intensity. All reactions were performed in triplicate.

### In vitro radioactive kinase assays

Assays were conducted by incubating 0.1 and 1 μM SidJ in 1X Protein Kinase buffer (NEB), with 10 mM CaCl_2_, 3 μM CaM, and 0.1 μg/μL MBP, and 1 μM SdeA 1-910. To initiate reactions an ATP mixture containing 100 μM cold ATP with 2.5 μCi ATPγP_32_ (Perkin Elmer) for 30 min at 30°C. Samples were then boiled and electrophoresed with SDS-PAGE gels which were dried and exposed for 2 hours to multiple days to visualize radioactive signal.

### In vitro AMPylation assays

SidJ and point mutants at a concentration of 0.5 μM were incubated with 50 mM Tris pH 7.5, 50 mM NaCl, with 5 μM CaM, 5 mM MgCl_2_, and in the presence or absence of 2 μM SdeA 231-1152 or SdeA truncations. Reactions were initiated with 2.5 μCi ATPαP_32_ for 30 minutes at 37°C. Samples were electrophoresed by SDS-PAGE, gels were dried and exposed between 1 hour and overnight to identify radioactive signals.

### *Legionella* strains and infections

*Legionella* strains used in this study include the wild type LP02 and the Dot/Icm deficient LP03 stains. The Δ*sidJ* strain was generated with triparental mating of the recipient WT strain, the pHelper strain and the donor E. coli DH5α λpir carrying the suicide plasmid pSR47s-*sidJ*. Integration of the plasmid was selected first with CYET plates containing 20 μg/mL of Kanamycin and then counterselected with CYET plates containing 5% sucrose. Colonies with genomic deletion were confirmed by PCR. Complementation strains were produced by electroporation of pZL507 plasmids containing SidJ wild type or D542A mutant into the Δ*sidJ* strain.

HEK293T cells were transfected with FCγRII and 4xFlag-Rab33 for 24 h. Bacteria of indicated Legionella strains were mixed with rabbit anti-legionella antibodies (1:500) at 37°C for 20 min. Cells were then infected with *L. pneumophila* strains at an MOI of 10 for 2 hours.

## Acknowledgement

This work is supported by National Institute of Health grant 5R01GM116964 (Y.M). This research used resources AMX 17-ID-1 of the National Synchrotron Light Source II, a U.S. Department of Energy (DOE) Office of Science User Facility operated for the DOE Office of Science by Brookhaven National Laboratory under Contract No. DE-SC0012704. The Life Science Biomedical Technology Research is primarily supported by the National Institute of Health, National Institute of General Medical Sciences (NIGMS) through a Biomedical Technology Research Resource P41 grant (P41GM111244), and by the DOE Office of Biological and Environmental Research (KP1605010). This investigation was supported by the National Institutes of Health under Ruth L. Kirschstein National Research Service Award (6T32GM008267) from the National Institute of General Medical Sciences (to MEM).

## Author contributions

Alan Sulpizio, Marena Minelli, Min Wan and Yuxin Mao conceived the general ideas for this work, performed the experiments, and wrote the manuscript. Xiaochun Wu, Paul Burrows, Ethan Sanford, Jung-Ho Shin, Byron Williams, Michael Goldberg, and Marcus Smolka contributed ideas and performed experiments.

---

**Figure 1—figure supplement 1.**
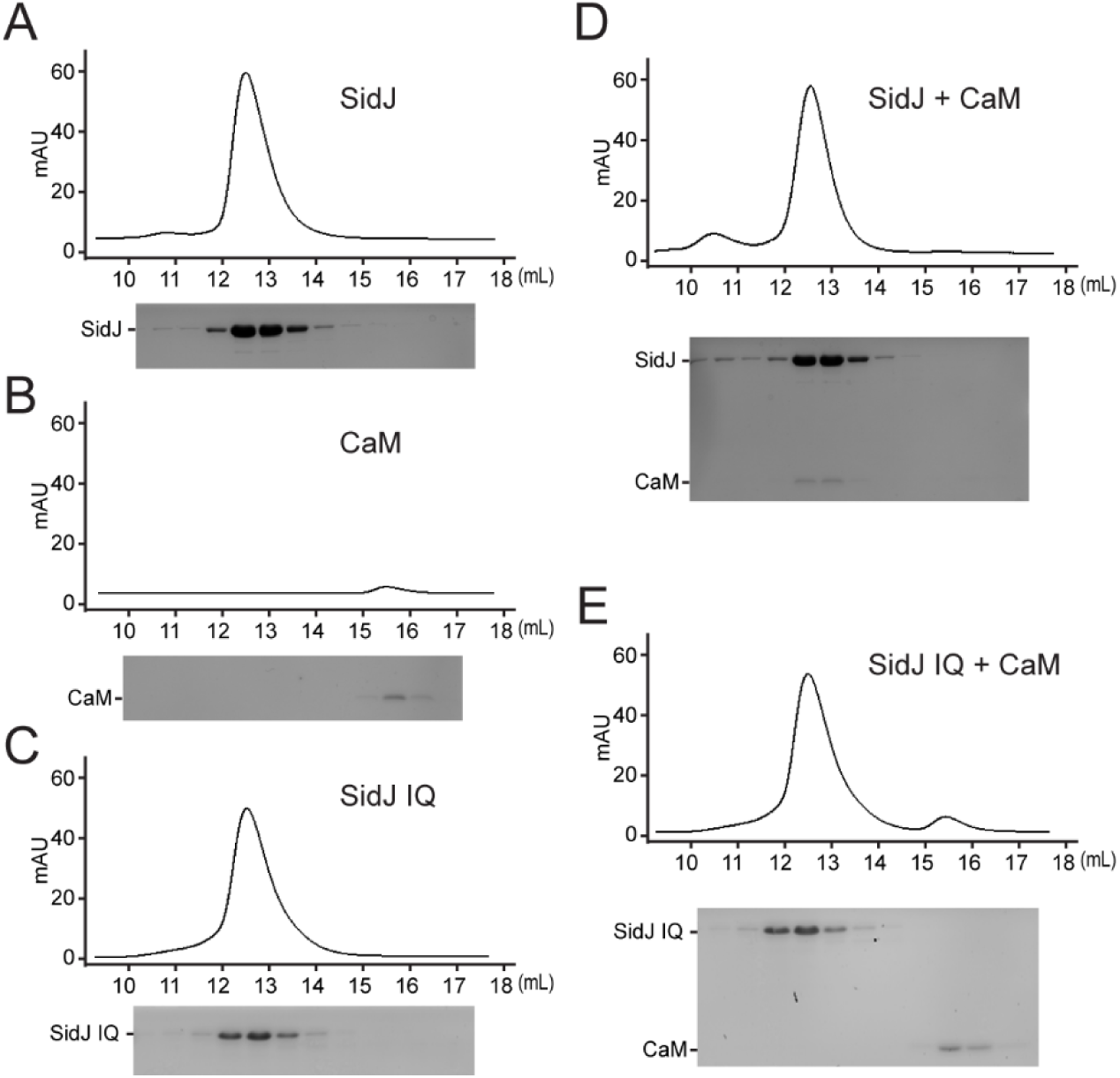
Size exclusion chromatography analysis of SidJ and CaM complex. (A-E) Size exclusion chromatogram profile of purified recombinant protein (top) and the peak fractions were visualized by SDS-PAGE followed by Coomassie staining (bottom). (A) SidJ; (B) CaM; (C) SidJ IQ mutant; (D) SidJ and CaM; and (E) SidJ IQ mutant and CaM.

**Figure 2—figure supplement 1.**
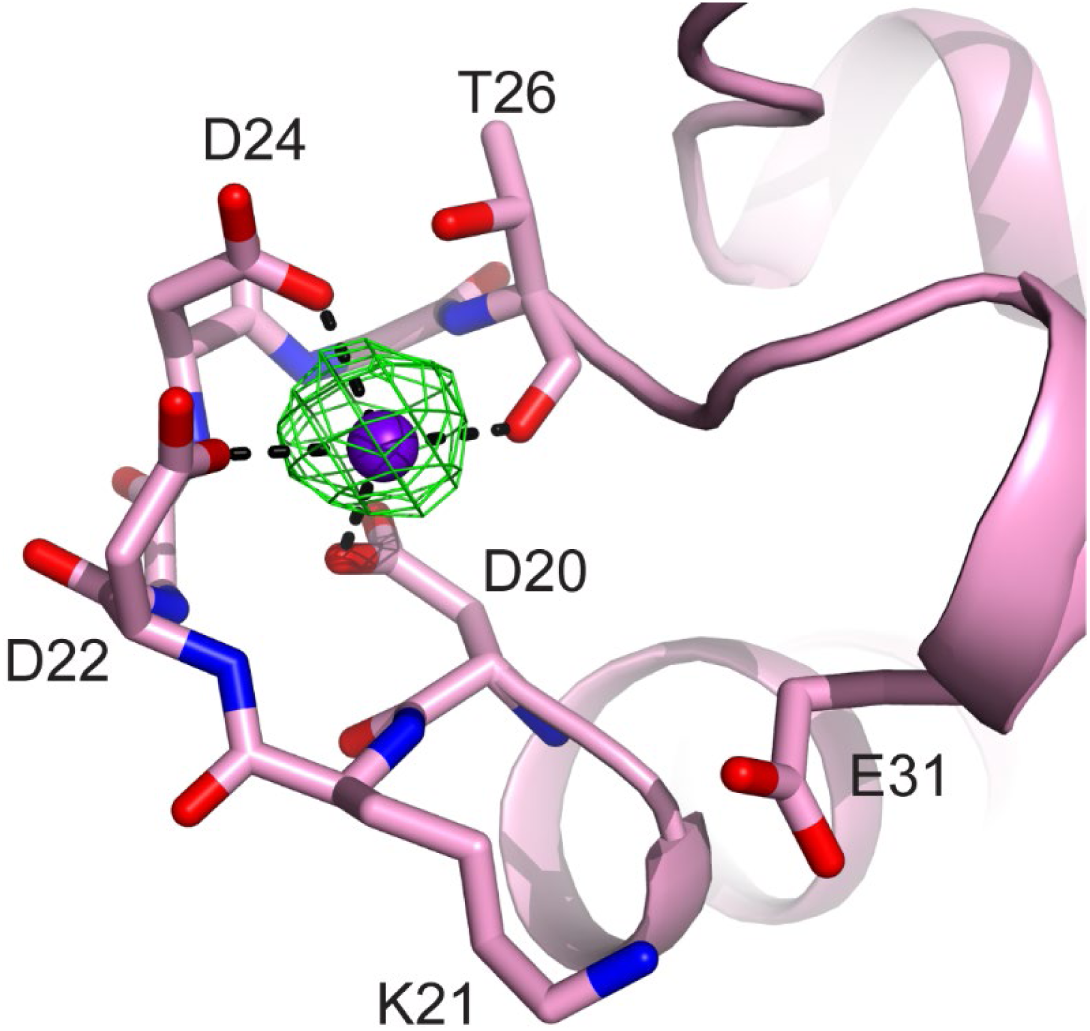
CaM EF-hand coordinated with one Ca^2+^. CaM is represented as pink cartoon with the residues that coordinate with the Ca^2+^ ion are shown as sticks. Green mesh represents F_o_-F_c_ difference map contoured to 3σ. Note that the conserved residue E31, which is responsible for chelation at the -Z coordination position in Ca^2+^ fully chelated CaM is shifted away from the Ca^2+^, indicating a weak Ca^2+^ binding to the CaM in the SidJ-CaM complex.

**Figure 3—figure supplement 1.**
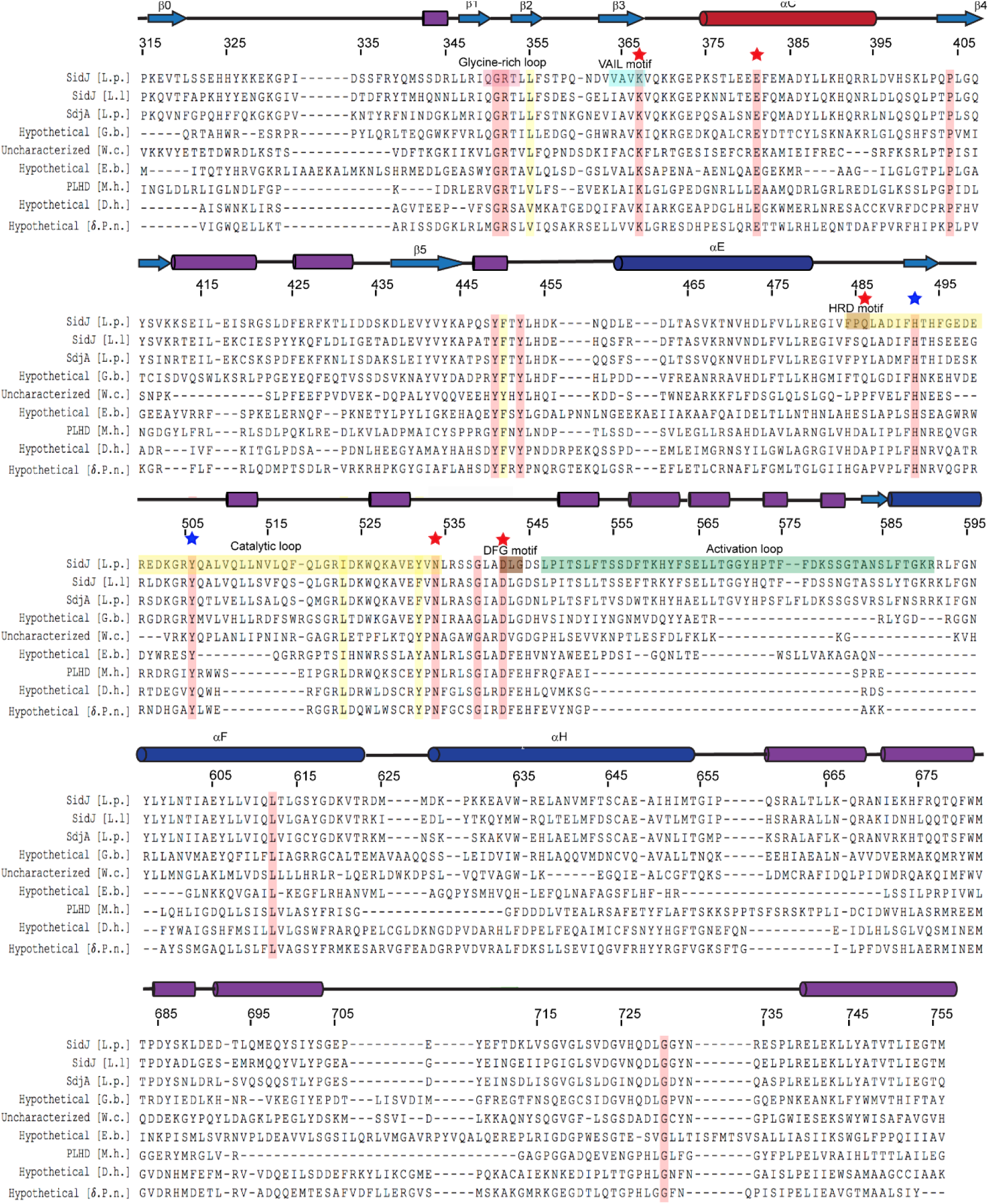
Multiple sequence alignment of SidJ kinase-like domain homologues. The NCBI BLAST server was used to identify homologous proteins to SidJ. Sequences corresponding to the kinase-like domain of SidJ (315-756) were aligned by Clustal Omega and colored using the MultAlin server (http://www.bioinformatics.org/sms/index.html). SidJ residue numbers are marked above the alignment with secondary structural elements drawn above. Identical residues are highlighted in red and similar residues in yellow. Kinase catalytic residues located in the active site are marked with red stars, while residues at the migrated-nucleotide binding pocket are marked with blue stars. Conserved kinase motifs are highlighted as follows: glycine-rich loop (red), VAIK motif (blue), HRD motif (gold), catalytic loop (yellow), DFG motif (brown) and activation loop (green). NCBI Accession numbers are as follows: SidJ *Legionella pneumophila*, AAU28221; SidJ *Legionella longbeachae*, RZV23241; SdjA *Legionella pneumophila*, AAU28568; hypothetical protein A3E83_09250, *Gammaproteobacteria bacterium RIFCSPHIGHO2_12_Full_41_20,* OGT46295.1; Putative uncharacterized protein*, Waddlia chondrophila 2032/99,* CCB91008.1; Hypothetical protein COB53_07685*, Elusimicrobia bacterium,* PCI37048.1; PBS lyase HEAT domain protein repeat-containing protein*, Methanosaeta harundinacea* KUK97762.1; Hypothetical protein*, Desulfovibrio hydrothermalis* WP_015335088.1; Hypothetical protein*, Delta proteobacterium NaphS2*, WP_006420030.1.

**Figure 3—figure supplement 2.**
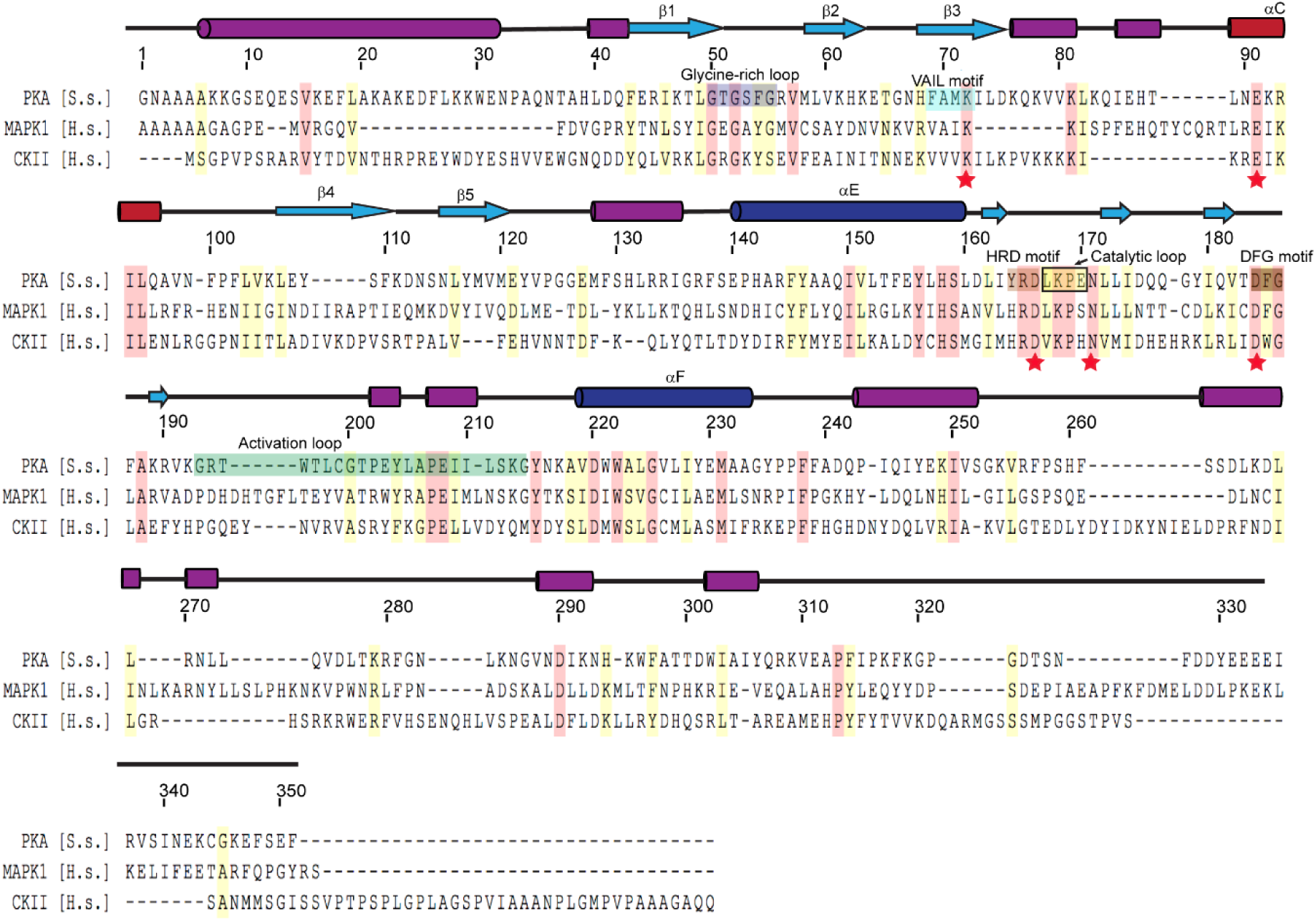
Multiple sequence alignment of representative protein kinases. The secondary structure of PKA is labeled above the sequence, and PKA residue numbers are marked on the top of the alignment. Identical residues are highlighted in red and similar residues in yellow. Kinase catalytic residues are marked with red stars and conserved kinase signature motifs are highlighted in a similar color scheme to SidJ kinase-like domain. Note that the catalytic loop in protein kinase contains only 4 amino acids, while the catalytic loop of SidJ is comprised of a large insertion (> 40 amino acids, Figure 3-Figure supplement 1). Accession numbers are as follows: cAMP-dependent protein kinase catalytic subunit alpha, *Sus scrofa,* P36887.4; Mitogen-activated protein kinase, *Homo sapiens,* P28482; Casein kinase II subunit alpha, *Homo sapiens*, P68400.

**Figure 3—figure supplement 3.**
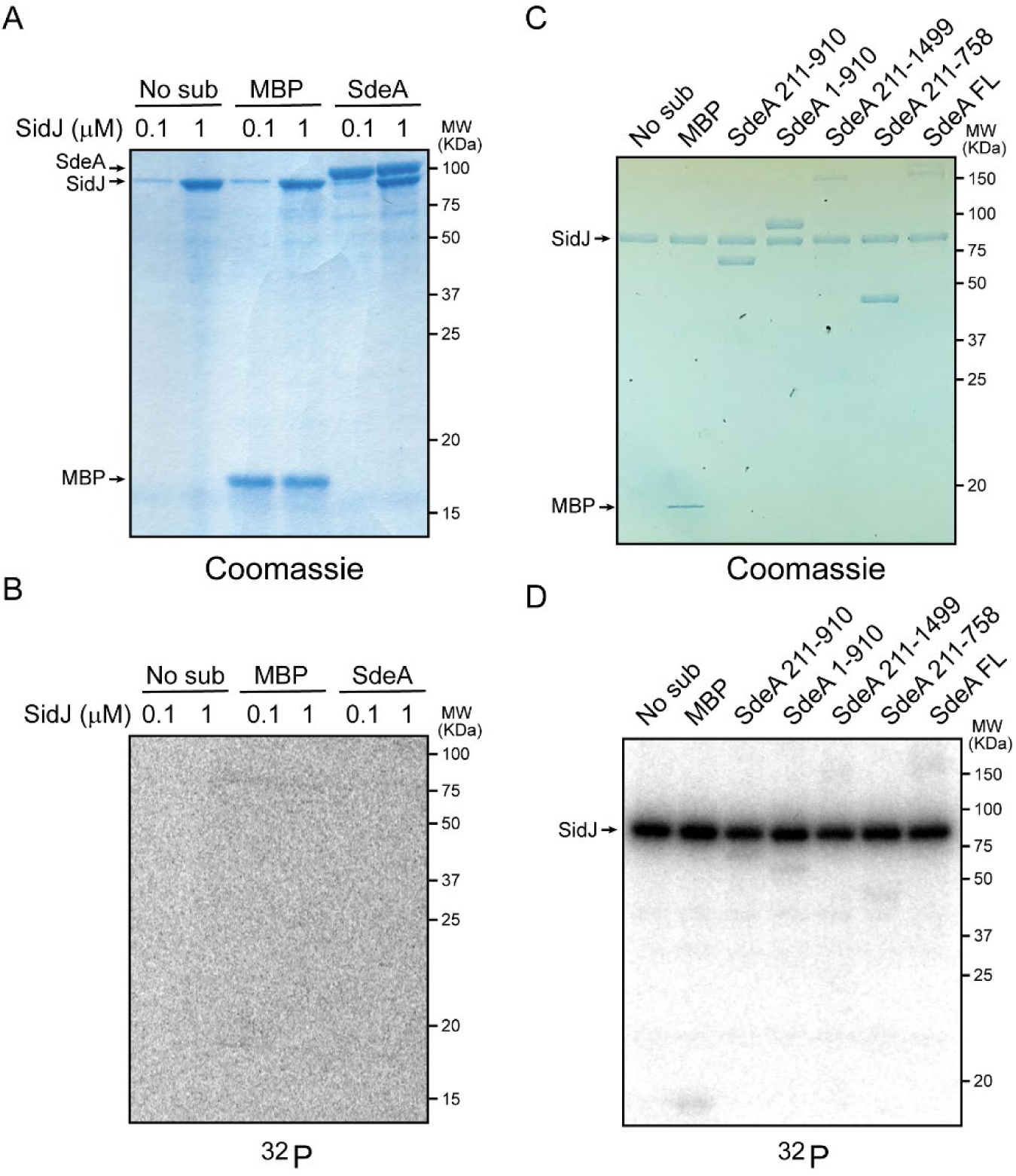
SidJ lacks canonical kinase activity but exhibits auto-AMPylation activity. (A) Two concentrations of SidJ 0.1 μM and 1 μM were incubated with CaM, MgCl_2_ and [γ-^32^P]ATP without substrate, with MBP, and with SdeA 1-910 for 30 minutes at 37°C. Proteins were separated by 12% SDS-PAGE and visualized with Coomassie stain. (B) Autoradiogram of gel shown in (A). Exposure time: 2 hours. (C) SidJ was incubated with CaM, MgCl_2_ and [α-^32^P]ATP without substrate, with MBP, and indicated recombinant SdeA truncations. Proteins were separated by 8% SDS-PAGE and visualized with Coomassie stain. (D) Autoradiogram of gel shown in (C). Exposure time: 1 hour. The bands corresponding to SidJ showed strong radiographic signals, indicating auto-AMPylation of SidJ.

**Figure 4—figure supplement 1.**
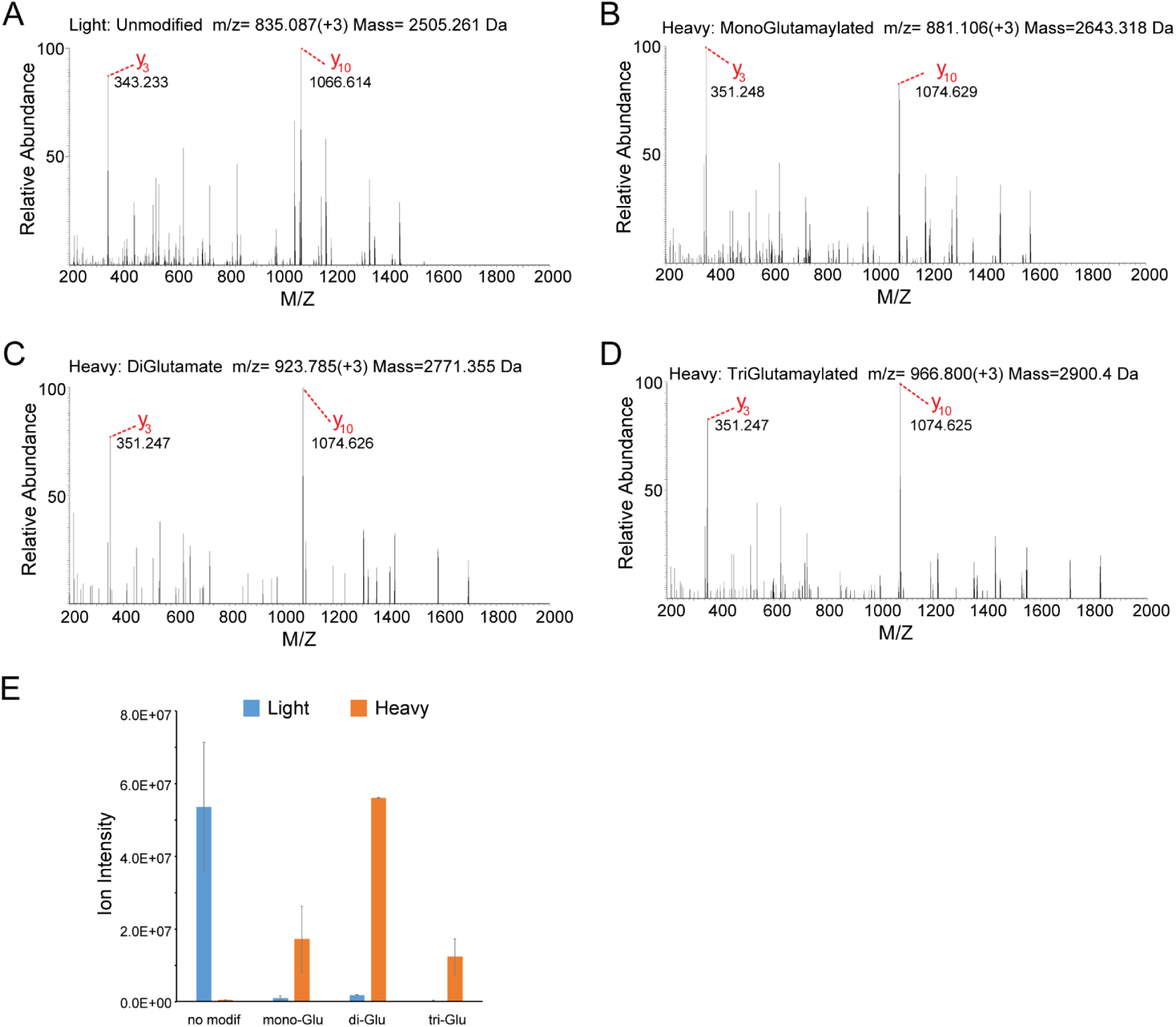
MS/MS analysis of the SdeA peptide modified by SidJ. (A) The MS2 spectrum of the SdeA peptide (residues 855-877, prepared from light medium) displays two signature ions, Y3 and Y10 ions, which correspond to the two Y ions generated from two labile prolines in the peptide. (B) A SdeA peptide (prepared from heavy medium) exhibits the same two signature ions (Y3 and Y10) but with a mass increase of 129.043 Da after subtraction of 8.014 Da corresponding to the heavy Lys and 1 Da corresponding to natural ^13^C incorporation in the peptide. The 129.043 Da corresponds to the addition of a glutamate residue. (C) A similar SdeA peptide (prepared from heavy medium) produces Y3 and Y10 ions, but with a mass increase of 2 x 129.043 Da after accounting for the heavy Lys. (D) A SdeA peptide (prepared from heavy medium) exhibits the same two signature ions (Y3 and Y10) but with a mass increase of 3 x 129.046 Da. (E) Quantification of ion intensity for heavy and light samples of unmodified, mono-glutamylated, di-glutamylated, and tri-glutamylated SdeA mART peptides. Data are shown as means ± STD of three independent mass spectrometry data collections.

**Figure 4—figure supplement 2.**
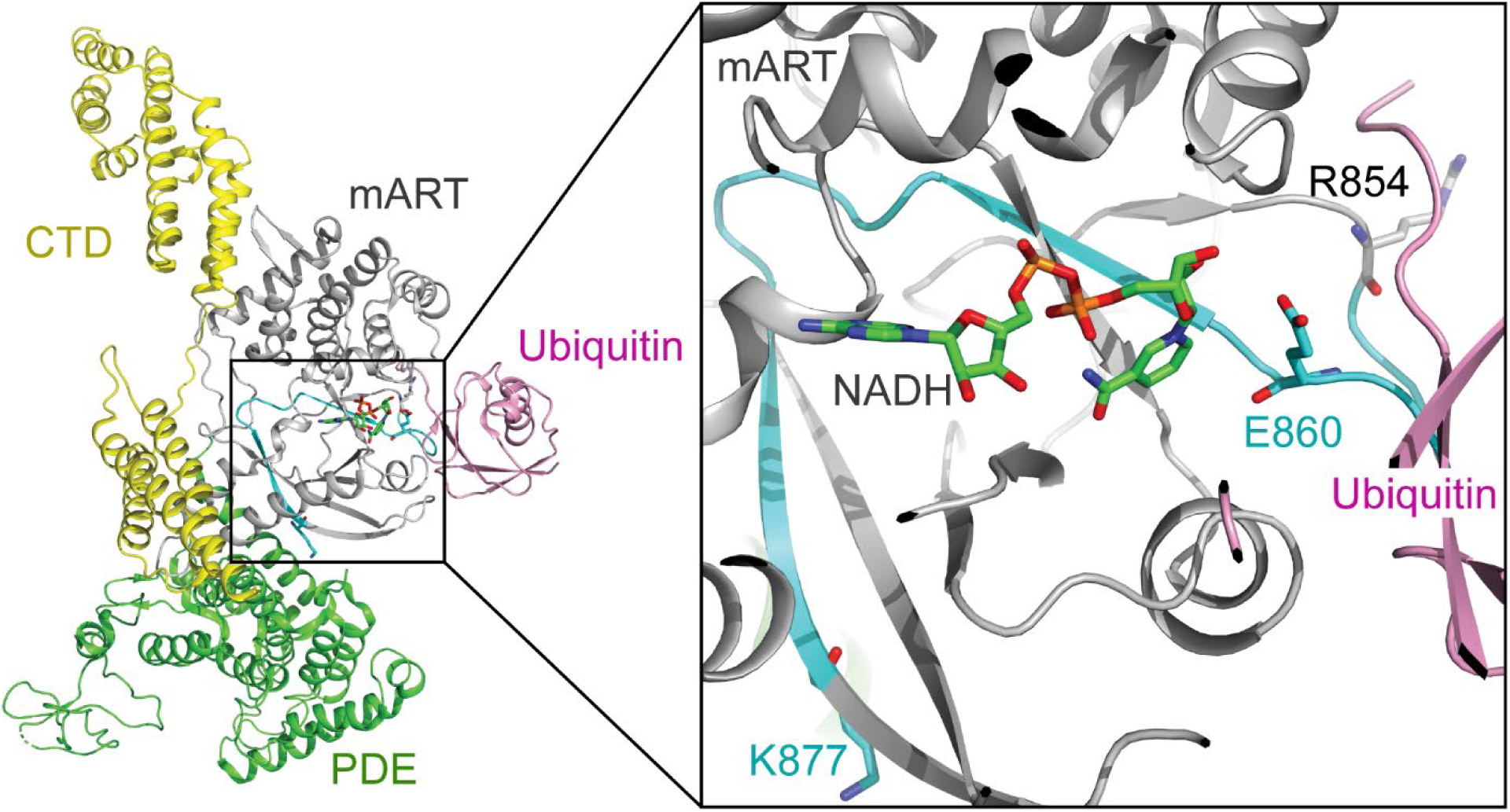
The structural context of the SdeA peptide modified by SidJ. Left: Overall structure of SdeA in complex with Ub and NADH (PDB ID: 5YIJ). Right: enlarged view of the mART active site of SdeA. The peptide (residues 855-877) shown in cyan was polyglutamylated at residue E860 (shown in sticks) by SidJ as detected by MS/MS analysis. The NADH displayed in sticks and colored in green.

**Figure 6—figure supplement 1.**
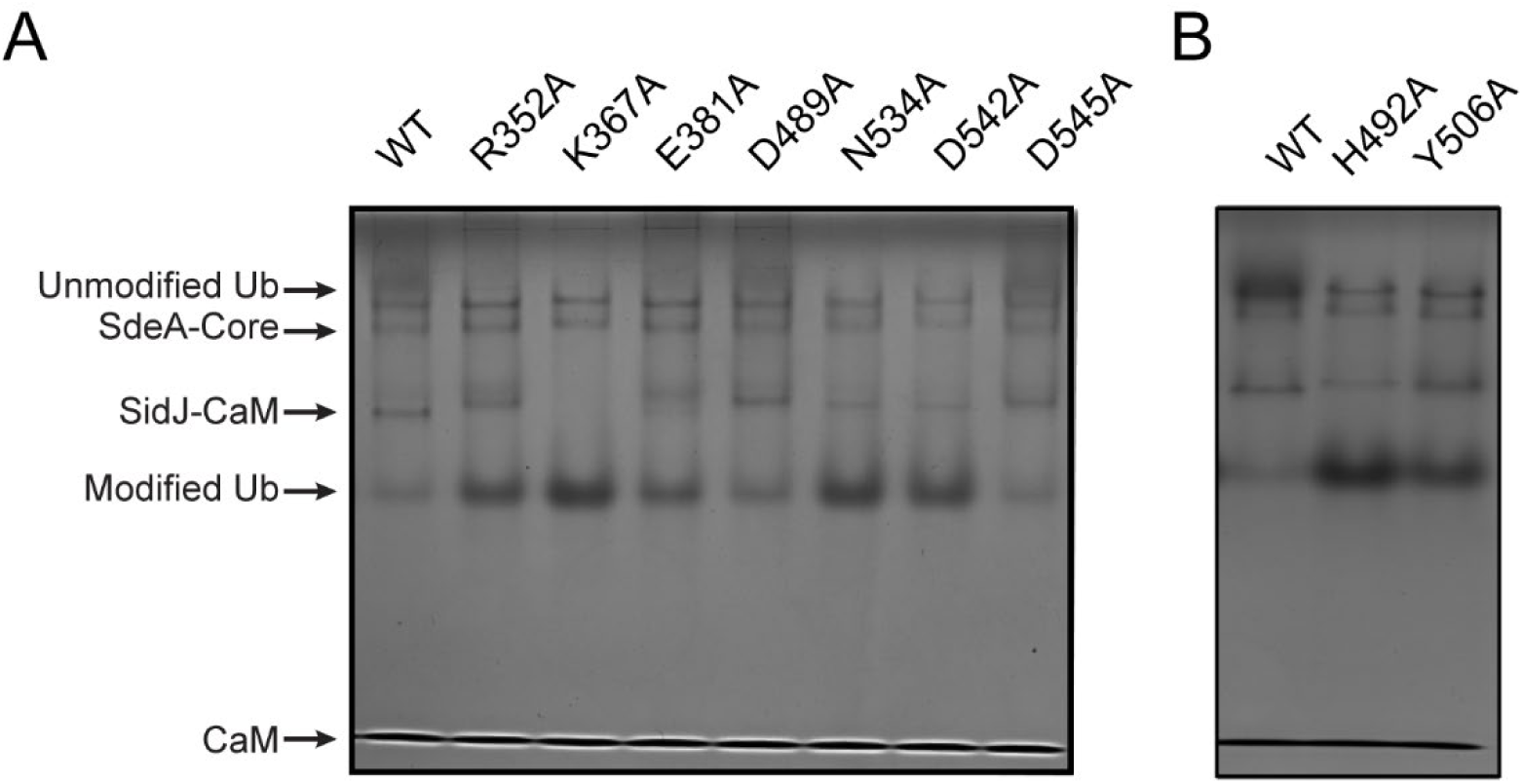
Inhibition of SdeA-catalyzed Ub ADP-ribosylation by SidJ mutants. (A) SdeA core was first treated by SidJ or its kinase active site mutants for 30 minutes at 37 °C. After the treatment, the SdeA mediated ADP-ribosylation of Ub was initiated by addition of Ub and NAD^+^ to the reaction mixture and further incubated for 30 minutes at 37°C. Reaction products were analyzed by Native PAGE and visualized with Coomassie stain. (B) The inhibition of SdeA-catalyzed ADP-ribosylation of Ub by SidJ nucleotide binding pocket mutants. The experiments were performed as in A.

**Figure 7—figure supplement 1.**
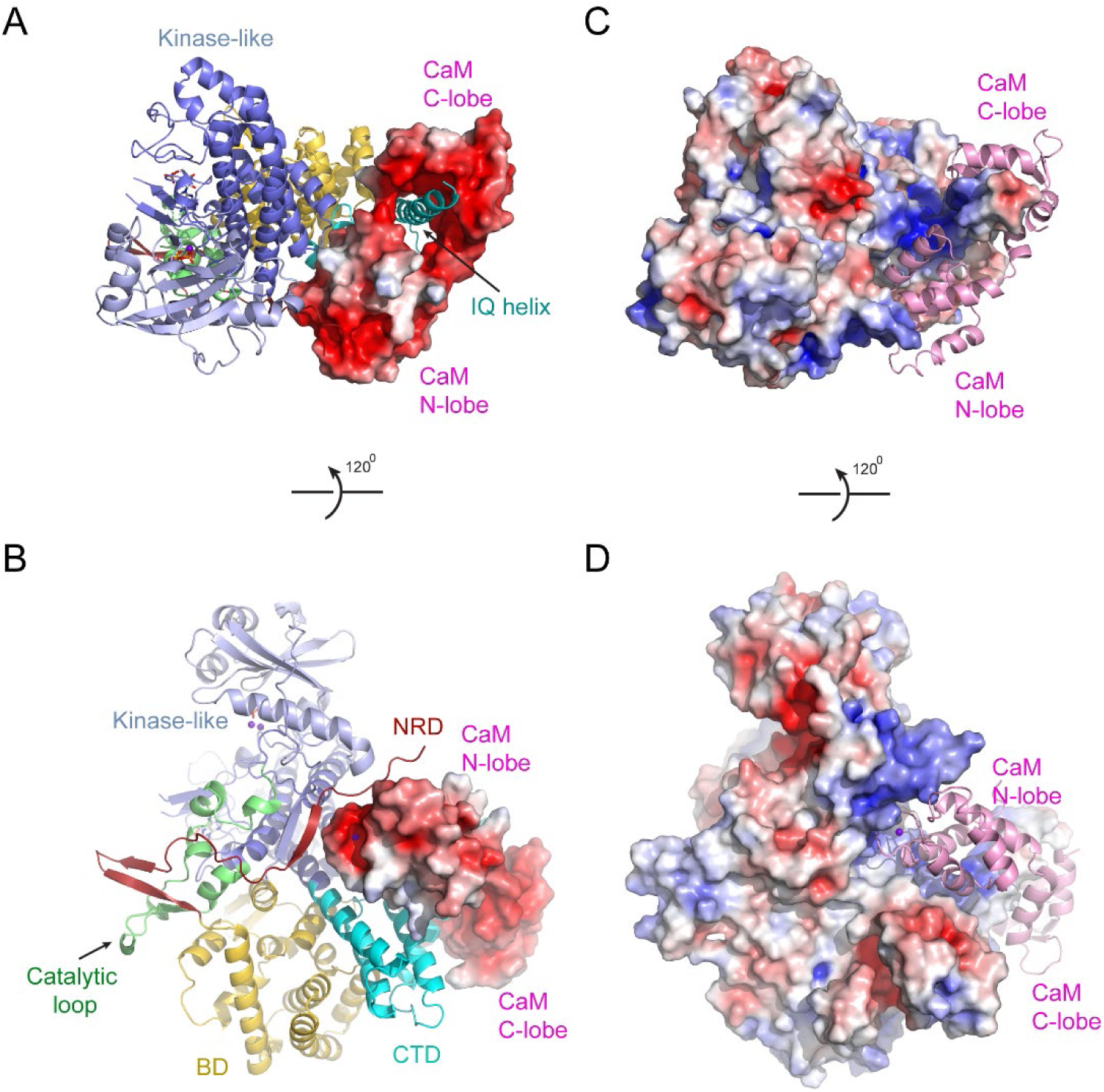
Electrostatic surface potential analysis of the interaction between SidJ and CaM. (A) The SidJ-CaM complex with SidJ depicted in ribbon and CaM shown with a surface representation colored by electrostatic surface potential with red being negatively charged with -5 eV and charged blue being positively charged with +5 eV. (B) A 120° rotated view of structure in (A). CaM is highly negatively charged as shown from both views. (C) The SidJ-CaM complex with SidJ presented in surface, which is colored according to its electrostatic surface potential (red: -5 eV; blue: +5 eV) and CaM in ribbon. (D) A 120° rotated view of structure in (C). The regions of SidJ interfaced with CaM are significant positively charged as shown from both views.

**Figure 7—figure supplement 2.**
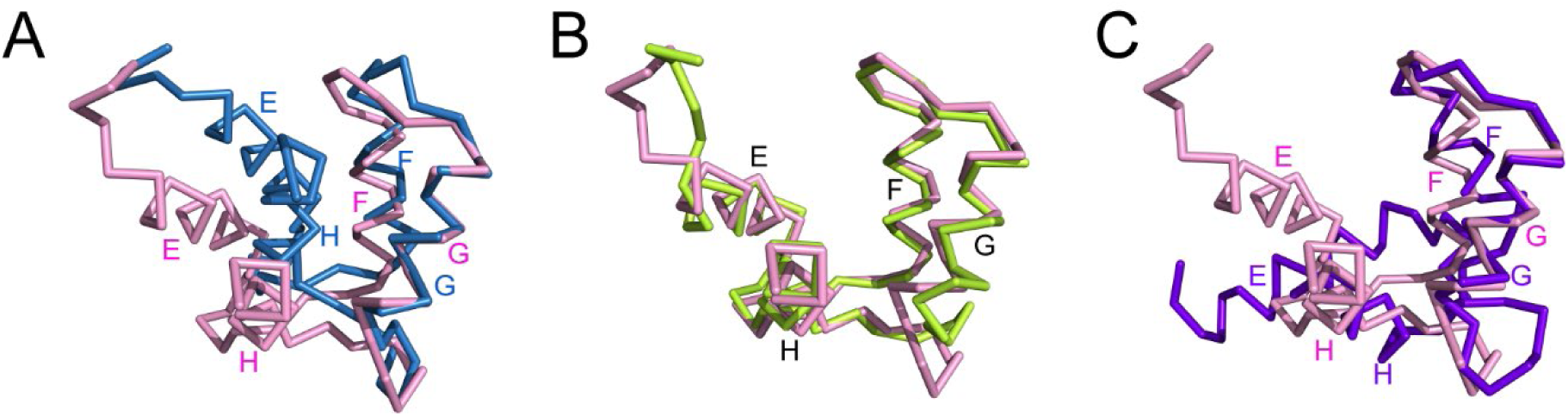
The C-lobe of CaM in the SidJ-CaM complex adopts a semi-open conformation. (A) A structural comparison of the CaM C-lobe in SidJ-CaM complex (pink) with that in apo-CaM (blue) (PDB ID: 1CFC). (B) Structural overlay of the CaM C-lobe in SidJ-CaM complex (pink) with CaM C-lobe (green) bound to the first IQ-motif in myosin V-CaM complex (PDB ID: 2IX7). (C) A structural comparison of the CaM C-lobe in SidJ-CaM complex (pink) with that (purple) in Ca^2+^ fully chelated CaM (PDB ID: 1CDL). The structures are overlaid in reference to helices F and G.

**Figure 7—figure supplement 3.**
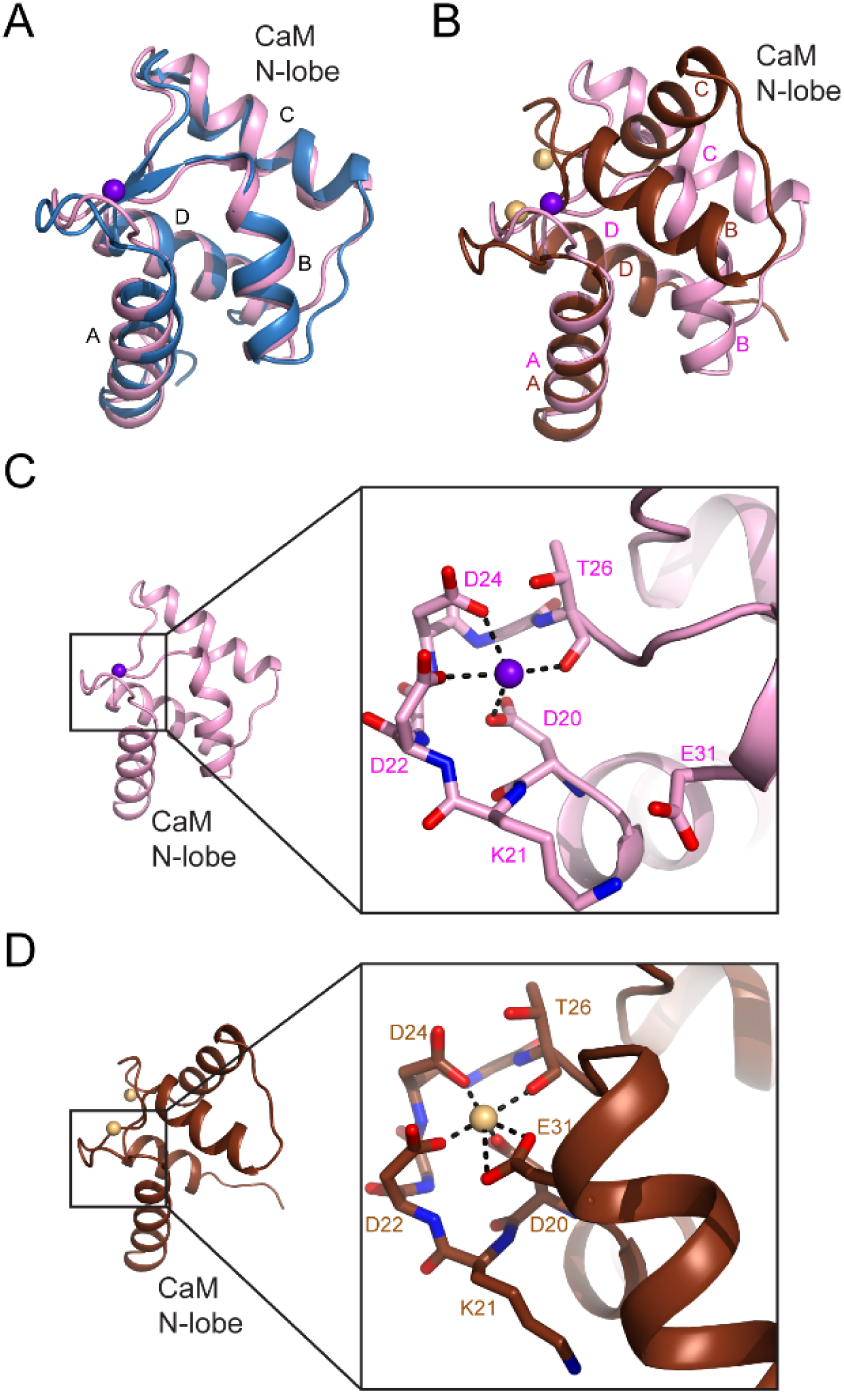
The N-lobe of the CaM in the SidJ-CaM complex adopts a closed conformation. (A) A structural comparison of the CaM N-lobe in the SidJ-CaM complex (pink) with that in apo-CaM (blue) (PDB ID: 1CFC). (B) A structural comparison of the CaM N-lobe in the SidJ-CaM complex (pink) with that in Ca^2+^ chelated CaM (brown) (PDB ID: 1CDL). (C) The CaM N-lobe in the SidJ-CaM complex is weakly bound by a Ca^2+^ ion as it is missing the –Z coordination due to the position of E31 away from the ion binding site. Thus, the N-lobe maintains a closed conformation. (D) Ca^2+^ chelating at the same binding site in (C) for Ca^2+^ saturated CaM (PDB ID: 1CDL). E31 is fully engaged in coordination with the bound Ca^2+^ ion and is responsible for the open conformation of the N-lobe.

**Figure 7—figure supplement 4.**
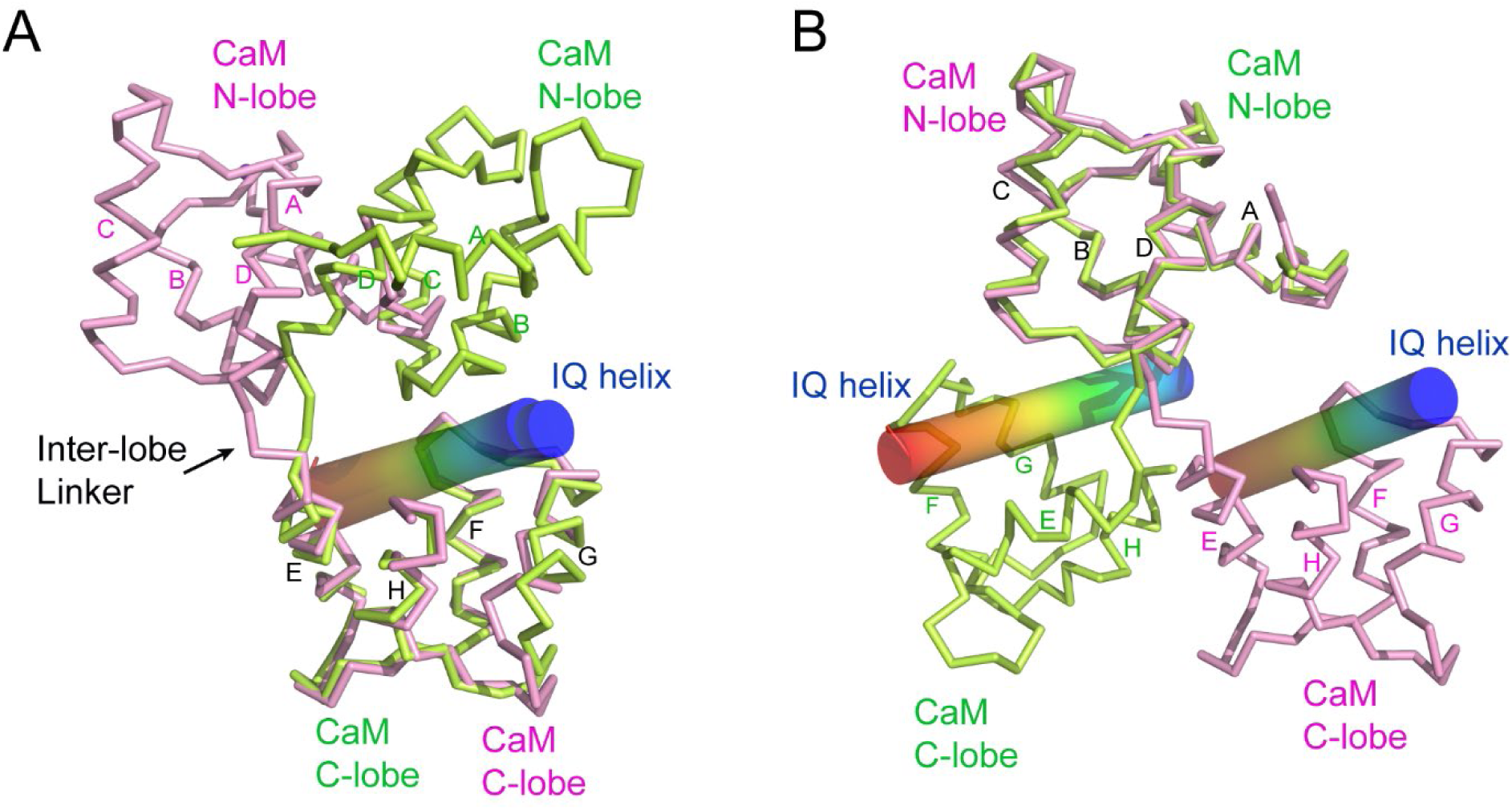
CaM adopts a unique conformation in the SidJ-CaM complex. (A) Structural comparison of CaM (pink) bound to SidJ IQ helix with that (green) bound to myosin V IQ1 (PDB ID: 2IX7). IQ helixes displayed as cylinders colored by spectrum from blue to red from the N-terminal to C-terminal ends, respectively. CaM structures are aligned respective to their C-lobes. (B) Structural overlay of same two structures shown in (A) with CaM structure aligned in reference to their N-lobes.

